# CatIF-RL: Activity-Oriented Enzyme Sequence Design by Steered Inverse Protein Folding

**DOI:** 10.64898/2026.05.11.724288

**Authors:** Yanheng Li, Jialong Xiong, Yuxin Zhang, Tong Cai, Chuan Fu, Shutong Li, Wei Xu, Ruoyi Lyu, Zhaoyang Chen, Zheng Guo, Xinqi Gong, Feng Wang

## Abstract

Protein inverse folding models are designed to generate amino acid sequences compatible with a given backbone structure, but they are not explicitly optimized for specific biological functions. Here, we present CatIF-RL, a framework that steers a graph-based denoising diffusion inverse folding model toward designing enzyme variants with enhanced catalytic activity. CatIF-RL first adapts the inverse folding model to enzyme structural data, then introduces activity-oriented preference signals using predicted catalytic constant (*k*_cat_) as the optimization objective, enabling specialization through generative dataset curation and group-relative policy optimization (GRPO). This process iteratively shifts the sequence distribution toward higher predicted *k*_cat_ while constraining sequence divergence to sequences that remain compatible with the input structure. On the independent benchmark, CatIF-RL achieves an approximately four-fold increase in predicted *k*_cat_ relative to native enzymes, substantially outperforming representative inverse folding methods, while maintaining sequence recovery (0.55) and structural fidelity, and supporting motif-preserving partial sequence design. CatIF-RL establishes a practical framework for activity-oriented enzyme design and provides a generalizable strategy for steering structure-conditioned protein generation toward functional optimization.

## 1. Introduction

Enzymes hold irreplaceable value in industrial biomanufacturing, pharmaceutical synthesis, and green chemistry, where their catalytic activity, substrate specificity, and stability directly dictate process performance^1–3^. Yet the properties selected through natural evolution are tuned to physiological contexts rather than to the demands of industrial or biotechnological applications. Engineering enzymes with enhanced or tailored catalytic performance has therefore become a fundamental challenge in protein design^4,5^. Classical campaigns based on directed evolution have produced numerous successes^6–8^, but their reliance on heuristic mutagenesis at a few residue positions inherently limits the sequence space they can traverse and makes them dependent on high-throughput screening and iterative selection, processes that are costly and time-consuming^9^. To overcome these bottlenecks, machine-learning-driven computational protein design has advanced rapidly over the past decade, giving rise to two complementary paradigms for enzyme engineering^10,11^. The first is the discriminative paradigm, in which models are trained to predict the effect of mutations on protein fitness or activity. Representative examples include the protein-language-model-based variant effect predictor ESM-1v^12^ and enzyme turnover number predictors such as DLKcat^13^, UniKP^14^, and CataPro^15^. These models can score candidate mutations with high throughput, substantially accelerating in silico evaluation of mutational libraries; however, they still depend on external search strategies (e.g., Monte Carlo sampling, genetic algorithms, or heuristic site enumeration) to traverse sequence space, and their efficiency is ultimately bounded by the number of oracle queries and the inductive biases of the search procedure itself^9,16,17^. The second is the generative paradigm, which performs conditional sampling directly at the level of the sequence distribution. This reframes sequence-space exploration entirely; models no longer passively score candidates but actively generate them under specified conditions, and can sample far beyond the neighborhood of naturally occurring homologs^18–21^. This shift has positioned generative models as a central direction for enzyme design and engineering.

Among the various branches of generative protein design, structure-conditioned generation is particularly well-suited to enzyme engineering^22,23^. Under this paradigm, a common decomposition divides the design problem into two complementary subproblems: (i) generating a protein backbone structure compatible with the desired function, and (ii) generating an amino acid sequence compatible with a given backbone. The latter is commonly referred to as protein inverse folding^24^. Inverse folding directly models the structure-to-sequence mapping without relying on natural homologs, enabling sampling that extends well beyond native sequence space. This yields a structurally constrained yet sequence-wise unconstrained design regime that aligns closely with the needs of enzyme engineering. Multiple representative models have emerged within this paradigm: ESM-IF transfers priors from large-scale protein language models to structure-conditioned sequence generation^25^; ProteinMPNN leverages graph neural networks to improve local structural consistency and design accuracy and has become the de facto baseline in the field^26^; PiFold introduces refined residue featurization and one-shot parallel decoding, improving efficiency and sequence diversity simultaneously^27^; LigandMPNN explicitly incorporates ligand and cofactor context into message passing, supporting binding-site-aware sequence design^28^; and the recently proposed ABACUS-T casts inverse folding as a discrete denoising diffusion process over sequence space and unifies ligand structures, multiple conformational states, and multiple sequence alignments (MSAs) within a single multimodal framework, thereby improving the precision of functional sequence generation^29^. Collectively, these methods have steadily advanced in structural representation and architectural design, progressively approaching the native distribution in both structural compatibility and sequence recovery, and have been successfully applied to the design of binders, antibodies, enzymes, and other functional proteins^30–34^.

Despite these advances, the training objectives of mainstream inverse-folding models remain centered on the negative log-likelihood of native sequences or on structural compatibility itself, without any explicit optimization of downstream biochemical properties such as catalytic activity^35,36^. This objective fundamentally biases models toward sampling from a native-like distribution, limiting their ability to explore functionally extreme or application-optimized regions of sequence space—an outcome at odds with the goals of enzyme engineering. Notably, reward-based post-training paradigms from adjacent fields, such as RLHF, DPO, and GFlowNet, have been used to steer generative models from distributional imitation toward objective-directed generation^37–40^. Introducing such paradigms into inverse-folding generation offers a path toward explicitly optimizing enzyme activity while preserving structural compatibility^35,41–43^.

To address this gap, we present CatIF-RL, a generalizable framework that recasts an inverse-folding model as an activity-directed sequence generator for enzyme design and engineering (Figure 1). Taking predicted enzyme turnover number (*k*_cat_) as the primary design objective, CatIF-RL combines generative dataset curation (GDC) with reinforcement learning (RL) fine-tuning to progressively reshape the sampling distribution toward regions of high catalytic activity. Specifically, we construct a structure dataset from BRENDA-derived enzyme data^44^ and train a graph-based inverse-folding model^45^ to capture both backbone conformation and residue-level physicochemical features. Using this model, we then generate candidate variants in batch on training-set backbones, filter them with an ensemble score derived from three independent catalytic activity predictors (DLKcat^13^, UniKP^14^, CataPro^15^) in conjunction with structural plausibility metrics, and train a new inverse-folding model from scratch on the resulting GDC dataset of high-predicted *k*_cat_ variants. Building on this checkpoint, CatIF, we further fine-tune the model using KL-regularized group-relative policy optimization (GRPO; KL refers to Kullback– Leibler divergence)^46^, which updates the sampling distribution toward higher predicted *k*_cat_ while preserving fold compatibility. In addition, we develop a subgraph inpainting sampling scheme tailored to the discrete-graph diffusion model, enabling localized sequence redesign under fixed catalytic residues or other functionally critical motifs, a setting common in enzyme engineering^47^. On a held-out benchmark with strict data-leakage control, CatIF-RL substantially outperforms representative inverse-folding baselines in predicted catalytic activity: its generated enzyme variants achieve, on average, approximately a four-fold increase in predicted *k*_cat_ relative to the corresponding native enzymes, while maintaining reasonable sequence recovery and predicted structural plausibility. These results demonstrate that the proposed post-training framework can effectively shift an inverse-folding model from native-distribution imitation to activity-directed generation, providing a generalizable training paradigm for functional optimization of structure-conditioned protein design, particularly for enzyme catalytic performance.

**Figure 1.**
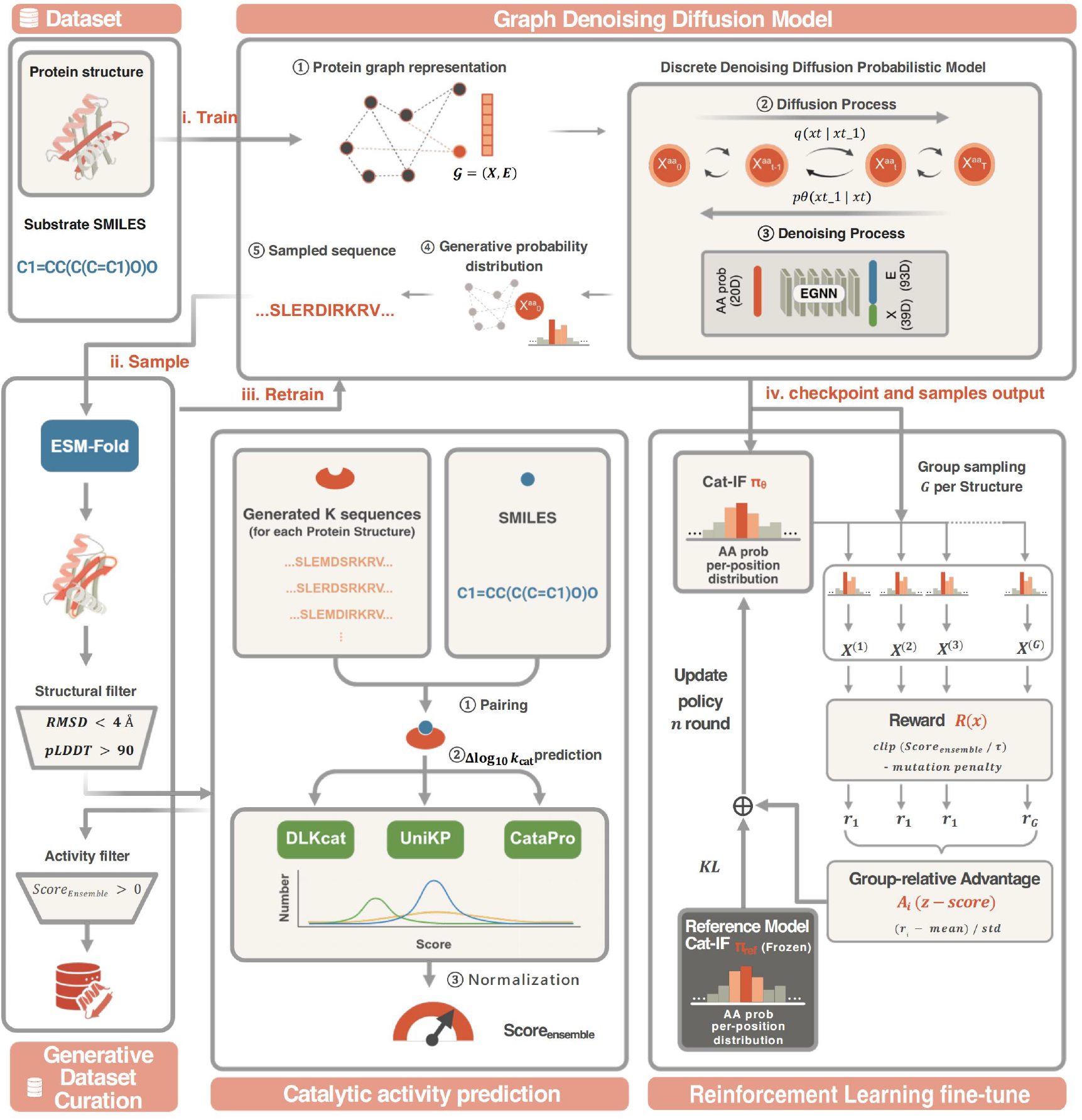
Overview of the CatIF-RL framework for activity-oriented enzyme sequence design. (i) The workflow begins with DLKcat-BRENDA enzyme sequences, substrate annotations, and predicted structures used to construct a leakage-controlled enzyme structure dataset. EnzymeIF is trained by adapting the GraDe-IF graph denoising diffusion inverse-folding architecture to enzyme structures with CATH-derived structural regularization. (ii) EnzymeIF is then used to generate candidate variants on training-set backbones. In GDC, candidates are first filtered by ESMFold-based structural plausibility and then scored by an ensemble of DLKcat, UniKP, and CataPro using normalized mutant-native activity differences. (iii) The activity-positive GDC variants are used to train CatIF from scratch. (iv) CatIF is subsequently fine-tuned by KL-regularized GRPO, producing CatIF-RL. Created in BioRender. Li, Y. (2026) https://BioRender.com/4spzumj

## 2. Methods

### 2.1 Dataset Construction and Leakage Control

We reconstructed a DLKcat-BRENDA enzyme kinetic dataset containing enzyme sequences, substrate SMILES strings, organism annotations, and associated catalytic measurements. The initial collection contained 16,839 organism–substrate–sequence records. After filtering sequences longer than 1,180 amino acids, removing sequences containing non-standard residues, and de-duplicating identical protein sequences, we obtained 7,713 distinct enzyme sequences. For each enzyme sequence, a structural model was predicted with ESMFold and used as the backbone condition for inverse-folding training and evaluation. The per-step filtering counts are detailed in Table S4, and the resulting sequence-length distribution is shown in Figure S1A.

To prevent leakage between model training and activity evaluation, we retained the original DLKcat split strategy^13^. The 1,423 sequences assigned to the DLKcat test split were held out as the independent benchmark set throughout the study. The remaining 6,290 sequences were used for model development and split into 5,661 training sequences and 629 validation sequences. In addition, to provide structural regularization during enzyme-domain adaptation, we incorporated CATH v4.2.0 general protein backbones into the training and validation sets. After preprocessing, 18,021 CATH structures were used for training and 608 for validation. These CATH structures were used only as general-protein structural regularizers and were not included in the held-out enzyme benchmark. The full per-source split is summarized in Table 1, with the corresponding leakage-control mechanics in Table S4.

**Table 1.** Dataset statistics. Processed enzyme structures were derived from the DLKcat-BRENDA dataset under the original DLKcat split. CATH v4.2.0 structures were incorporated as general-protein structural regularizers during EnzymeIF training.

| Data source | Train | Validation | Test | Total |
| --- | --- | --- | --- | --- |
| DLKcat-BRENDA | 5,661 | 629 | 1,423 | 7,713 |
| CATH v4.2.0 | 18,021 | 608 | - | 18,629 |
| Combined dataset | 23,682 | 1,237 | 1,423 | 26,342 |

### 2.2 GraDe-IF Backbone and EnzymeIF Training

CatIF-RL is built on the GraDe-IF architecture^45^, a graph denoising diffusion model for inverse protein folding (Figure 1). A protein backbone is represented as a residue graph 𝒢, where nodes *X* correspond to residues and edges *E* encode local spatial relationships. The model defines a discrete diffusion process over amino-acid node types and learns to reverse this corruption process conditioned on the backbone graph, residue-level geometric and physicochemical features, and secondary-structure information. The denoising network is implemented with an equivariant graph neural network (EGNN), and the forward transition kernel is based on a BLOSUM substitution matrix rather than a uniform categorical transition, thereby incorporating biologically informed amino-acid replacement priors into the diffusion process.

Using this architecture, we trained EnzymeIF as an enzyme-specialized structure-only inverse-folding model. EnzymeIF was trained on the DLKcat-BRENDA enzyme training structures together with the CATH-derived general protein structures described above. The enzyme structures provide the domain-specific training signal, whereas the CATH structures act as structural regularizers that broaden backbone geometry coverage during denoising training.

EnzymeIF used the same core architecture as GraDe-IF: an EGNN-based denoiser with 6 layers, hidden dimension 128, dropout 0.1, 500 diffusion steps, and a BLOSUM-based discrete transition kernel. Training minimized the residue-level cross-entropy between the predicted clean amino-acid distribution and the native sequence under the discrete diffusion objective. Unless otherwise specified, models were optimized with Adam using a learning rate of 5 × 10^−4^, weight decay 1 × 10^−5^, batch size 32, gradient clipping with maximum norm 1.0, and an exponential moving average decay of 0.995. Detailed training hyperparameters, checkpoint selection criteria, and training/validation curves are provided in Table S1 and Figure S2A.

### 2.3 Generative Dataset Curation

We used EnzymeIF to generate an activity-enriched training dataset through GDC (Figure 1). For each of the 6,290 enzyme backbones in the leakage-controlled training split, we sampled *K* = 10 sequences from the discrete diffusion reverse process, resulting in 62,900 candidate variants.

Candidate variants were filtered in two stages. First, structural plausibility was assessed by refolding each generated sequence with ESMFold and comparing the predicted structure with the corresponding reference backbone. Variants were retained if they satisfied backbone RMSD < <4 Å and mean pLDDT > 90, yielding 15,188 structure-valid variants. The per-candidate Avg pLDDT and backbone RMSD distributions used in this gate are shown in Figure S1B,C.

Second, structure-valid variants were evaluated with an ensemble of three pretrained catalytic turnover predictors: DLKcat, UniKP, and CataPro. For predictor *m*, each candidate sequence *x* paired with substrate *s* was scored relative to the corresponding native enzyme sequence *x*_native_: Predictor versions, repositories, and inference interfaces are listed in Table S5.

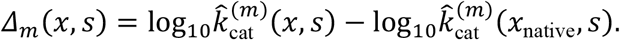

To place the three predictors on a comparable scale, we normalized each predictor-specific *Δ*_*m*_ distribution using robust quantile-range scaling:

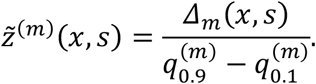

Here, 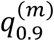 and 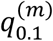 denote the 90th and 10th percentiles of the *Δ*_*m*_ distribution for predictor *m*, estimated over the 15,188 structure-valid candidate variants. This scaling adjusts for predictor-specific output ranges but does not subtract the median or otherwise re-center the candidate distribution.

The ensemble activity score was defined as the arithmetic mean of the three normalized predictor scores:

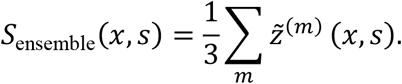

Variants with *S*_ensemble_(*x, s*) > 0 were retained as activity-positive candidates. This procedure produced 6,034 GDC variants, which were used for supervised training of CatIF. Pairwise inter-predictor agreement on this 25,970-triplet candidate pool is reported in Figure S4.

### 2.4 CatIF Supervised Training on the GDC Dataset

CatIF was trained from scratch on the 6,034 activity-positive variants obtained from GDC (Figure 1). CatIF used the same discrete diffusion inverse-folding architecture and training objective as EnzymeIF, but differed in the training distribution: whereas EnzymeIF was trained on enzyme structures plus CATH-derived structural regularizers, CatIF was trained only on the curated activity-positive GDC variant set. Thus, EnzymeIF served as the generator used to construct the GDC candidate pool, whereas CatIF was a separately trained activity-enriched inverse-folding model rather than a checkpoint continuation or distillation of EnzymeIF.

The CatIF model used an EGNN-based denoiser with 6 layers, hidden dimension 128, dropout 0.1, 500 diffusion steps, and a BLOSUM-based transition kernel. Training used Adam with learning rate 5 × 10^−4^, weight decay 1 × 10^−5^, batch size 32, gradient clipping with maximum norm 1.0, and exponential moving average decay 0.995. The full hyperparameter list, the lowest-validation-loss checkpoint selection (epoch 228), and the corresponding training/validation curves are documented in Table S2 and Figure S2B.

### 2.5 CatIF-RL Optimization by KL-Regularized GRPO

CatIF-RL further refines CatIF using KL-regularized GRPO. The policy *π*_*θ*_ is initialized from the supervised CatIF checkpoint, and the same CatIF checkpoint is used as the frozen reference policy *π*_ref_ for KL regularization. This design constrains the optimized policy to remain close to a fold-compatible inverse-folding generator while allowing the sampling distribution to shift toward higher predicted catalytic activity.

Training proceeds in outer rounds. In each round, the current policy generates candidate sequences for each backbone, and the candidates are scored using the activity predictor ensemble described above. Policy optimization is then performed on the pre-scored candidate set, without invoking the predictors during backpropagation. Unless otherwise specified, we used three outer rounds and two inner optimization epochs per round; the checkpoint after the second epoch of each round was used to generate candidates for the next round. The complete two-level loop (outer rounds + inner offline GRPO update) is given in Algorithm S2.

The activity component of the reward was derived from the ensemble score:

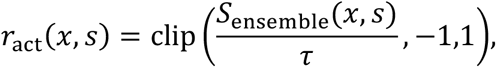

where *τ* is a scaling parameter controlling the sensitivity of the reward to ensemble-predicted improvement. To discourage excessive divergence from the native sequence, we added a mutation penalty based on the mutation fraction *μ*(*x, x*_native_):

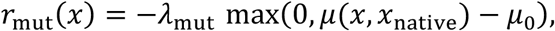

where *μ*_0_ is the free-mutation threshold and *λ*_*mut*_ is the penalty coefficient. The final reward was

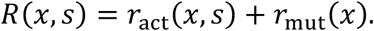

For each backbone, we sampled a group of *G candidate sequences* and computed group-relative advantages:

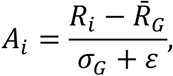

where 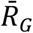 and *σ*_*G*_ denote the mean and standard deviation of rewards within the group. This group-relative normalization reduces sensitivity to imperfect global reward calibration and aligns the optimization procedure with the practical inverse-folding regime of generating multiple candidates per backbone.

The policy objective used length-normalized sequence log-probabilities to avoid length-induced bias. For sequence *x*_*i*_ of length *L*_*i*_, we define

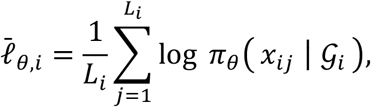

where 𝒢_*i*_ denotes the corresponding backbone graph. The KL-regularized GRPO objective was written as

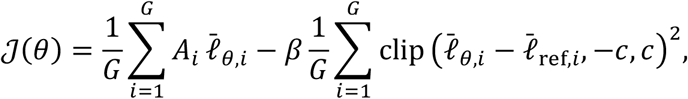

where 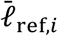 is the corresponding length-normalized log-probability under the frozen CatIF reference policy, *c* is the clipping range for the KL proxy, and *β* is the regular*i*zat*i*on coefficient. Full RL hyperparameters across the three rounds are provided in Table S3, and inner-loop convergence diagnostics (total loss, KL proxy, and reward × log-prob correlation) are shown in Figure S3.

### 2.6 Motif-Preserving Partial Design by Discrete Subgraph Inpainting

To support constrained enzyme redesign, we implemented a discrete subgraph inpainting sampler for motif-preserving partial sequence design. This setting is relevant when catalytic residues, metal-binding residues, or experimentally validated functional motifs must remain fixed while surrounding positions are redesigned.

Let *m* ∈ {0,1}^*L*^ be a binary mask over residues, where *m*_;_ = 1 denotes a fixed position and *m*_*i*_ = 0 denotes a designable position. Given a native sequence *x*_native_ and a backbone graph *𝒢*, fixed positions are enforced by sampling from the forward marginal of the native sequence at the corresponding diffusion step:

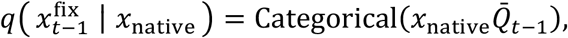

where 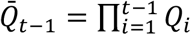 is the cumulative discrete transition matrix. This operation keeps fixed residues on the correct marginal trajectory of the forward diffusion process rather than allowing them to inherit uncertainty from model-predicted intermediate states.

For designable positions, the denoiser predicts a clean-sequence distribution and the reverse transition is obtained by marginalizing over the predicted clean amino-acid states:

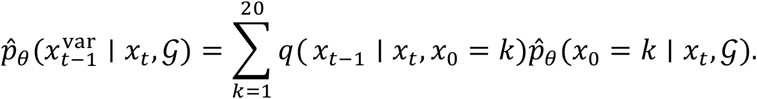

The fixed and designable regions are then fused residue-wise:

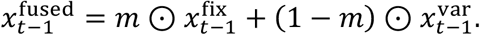

To improve compatibility at the boundary between fixed and designable regions, we further used inpainting-style resampling: the fused state was partially re-noised and re-denoised for a small number of cycles at each diffusion step.

### 2.7 Evaluation and Statistical Analysis

We evaluated CatIF-RL and representative inverse-folding baselines on the held-out DLKcat-aligned benchmark containing 1,423 enzymes. All functional evaluations were performed as paired mutant–native comparisons using the same protein and substrate annotations. For each test backbone, models were sampled in five independent rounds using five random seeds unless otherwise specified. The per-baseline sampling configuration (seed values, decoding temperature, sample count, ligand/MSA inputs, checkpoint version) is summarized in Table S6.

The primary functional metric was the protein-level change in predicted catalytic turnover:

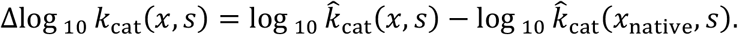

When a protein was associated with multiple substrates, Δlog_10_ *k*_cat_ was first averaged over substrates for that protein and then averaged across proteins. Reported values were averaged over the five sampling rounds.

Recovery Rate was defined as the fraction of positions in the generated sequence that matched the native sequence:

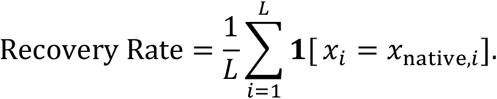

Recovery Rate was computed for each generated sequence, averaged over proteins within each round, and then averaged over the five sampling rounds.

Structural plausibility was evaluated by predicting the structure of generated sequences with ESMFold and comparing each predicted structure with the corresponding reference backbone. Backbone RMSD was used to measure global structural deviation, and mean pLDDT was used as a local confidence measure. Because refolding all five-seed outputs is computationally expensive, pLDDT, backbone RMSD, and SR were computed on a pre-specified structural-evaluation sample set generated under one random seed.

Success rate (SR) was defined as the fraction of generated variants satisfying a joint structure–function criterion:

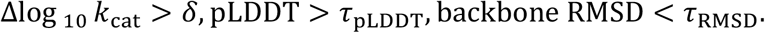

Unless otherwise specified, 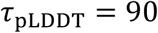 and *τ*_RMSD_ = 4Å. The main table reports SR at *δ* = 0 and *δ* = 1.0, corresponding to default activity improvement and a stringent one-log improvement threshold, respectively. Intermediate thresholds were used for additional threshold-sensitivity analyses where indicated.

When statistical comparisons were performed, they were conducted at the protein level. For Δlog_10_ *k*_cat_, Recovery Rate, pLDDT, and backbone RMSD, paired protein-level comparisons between CatIF-RL and each baseline were assessed using Wilcoxon signed-rank tests, with *p*-values adjusted by the Benjamini–Hochberg procedure. Protein-level bootstrap resampling with 10,000 resamples was used to estimate confidence intervals for mean differences. All statistical analyses were performed after aggregating replicate designs at the protein level to avoid treating multiple samples from the same backbone as independent observations.

## 3. Results

### 3.1 Domain Adaptation Establishes an Enzyme-Specialized Inverse Folding Prior

We first adapted the generic GraDe-IF inverse-folding architecture to the enzyme domain. The original GraDe-IF model provides a discrete diffusion inverse-folding reference trained for general protein structures, but it is not explicitly specialized for enzyme sequence–structure distributions. We therefore trained EnzymeIF on the DLKcat-BRENDA enzyme training set, with CATH-derived general protein backbones incorporated as structural regularizers.

On the DLKcat-aligned held-out benchmark, EnzymeIF substantially improved over the generic GraDe-IF reference across enzyme-relevant inverse-folding metrics. EnzymeIF achieved a Δlog_10_ *k*_cat_ of 0.2359, compared with 0.1115 for GraDe-IF. More importantly for the enzyme-domain adaptation stage, EnzymeIF increased Recovery Rate from 0.4617 to 0.7088, indicating a stronger ability to recover native-like enzyme sequences under the same backbone-conditioned generation setting. EnzymeIF also improved predicted structural plausibility, with pLDDT increasing from 75.73 to 83.74 and backbone RMSD decreasing from 5.3912 Å to 3.8566 Å. These results establish EnzymeIF as an enzyme-specialized structure-only generator and motivate its use as the candidate generator for the subsequent GDC stage.

(See Table 2 for the full benchmark and Table S7 for the corresponding paired Wilcoxon p-values and 95 % CIs.)

**Table 2.** Benchmark summary on the DLKcat-aligned held-out test set. Δlog_10_ *k*_cat_ is computed from paired mutant–native DLKcat predictions and averaged at the protein level. Recovery Rate is the fraction of residues matching the native sequence. pLDDT and backbone RMSD are computed using the pre-specified ESMFold structural-evaluation sample set. SR@*δ*=0 and SR@*δ*=1.0 report the fraction of variants satisfying the joint structural criteria and the corresponding minimum Δlog _10_ *k*_cat_ threshold.

| Model | $\Delta\log_{10}(k_{\text{cat}})$ | Recovery Rate | pLDDT | Backbone RMSD (Å) | $\text{SR}@ \delta=0$ | $\text{SR}@ \delta=1.0$ |
| --- | --- | --- | --- | --- | --- | --- |
| **CatIF-RL** | 0.6049 | 0.5533 | 77.87 | 5.3224 | 0.1266 | 0.0345 |
| **CatIF** | 0.3647 | 0.5719 | 79.16 | 4.8850 | 0.1145 | 0.0246 |
| **EnzymeIF** | 0.2359 | 0.7088 | 83.74 | 3.8566 | 0.1252 | 0.0274 |
| **GraDe-IF** | 0.1115 | 0.4617 | 75.73 | 5.3912 | 0.0141 | 0.0014 |
| **LigandMPNN** | -0.0550 | 0.5139 | 87.05 | 3.8881 | 0.1097 | 0.0338 |
| **ESM-IF** | 0.0751 | 0.5359 | 82.32 | 5.1993 | 0.0942 | 0.0274 |
| **ABACUS-T** | 0.0061 | 0.6042 | 85.67 | 4.0512 | 0.0654 | 0.0197 |
| **PiFold** | -0.0858 | 0.5298 | 85.47 | 4.2279 | 0.0893 | 0.0309 |
| **ProteinMPNN** | -0.1182 | 0.4704 | 86.11 | 3.8601 | 0.0696 | 0.0260 |

### 3.2 Generative Dataset Curation Constructs an Activity-Enriched Sequence Distribution

Using EnzymeIF, we generated 62,900 candidate variants from the 6,290 training-set enzyme backbones, with 10 sampled sequences per backbone. We then applied a two-stage GDC procedure to select variants that were both structurally plausible and activity-positive under the predictor ensemble. (Figure 2A summarises this funnel.)

**Figure 2.**
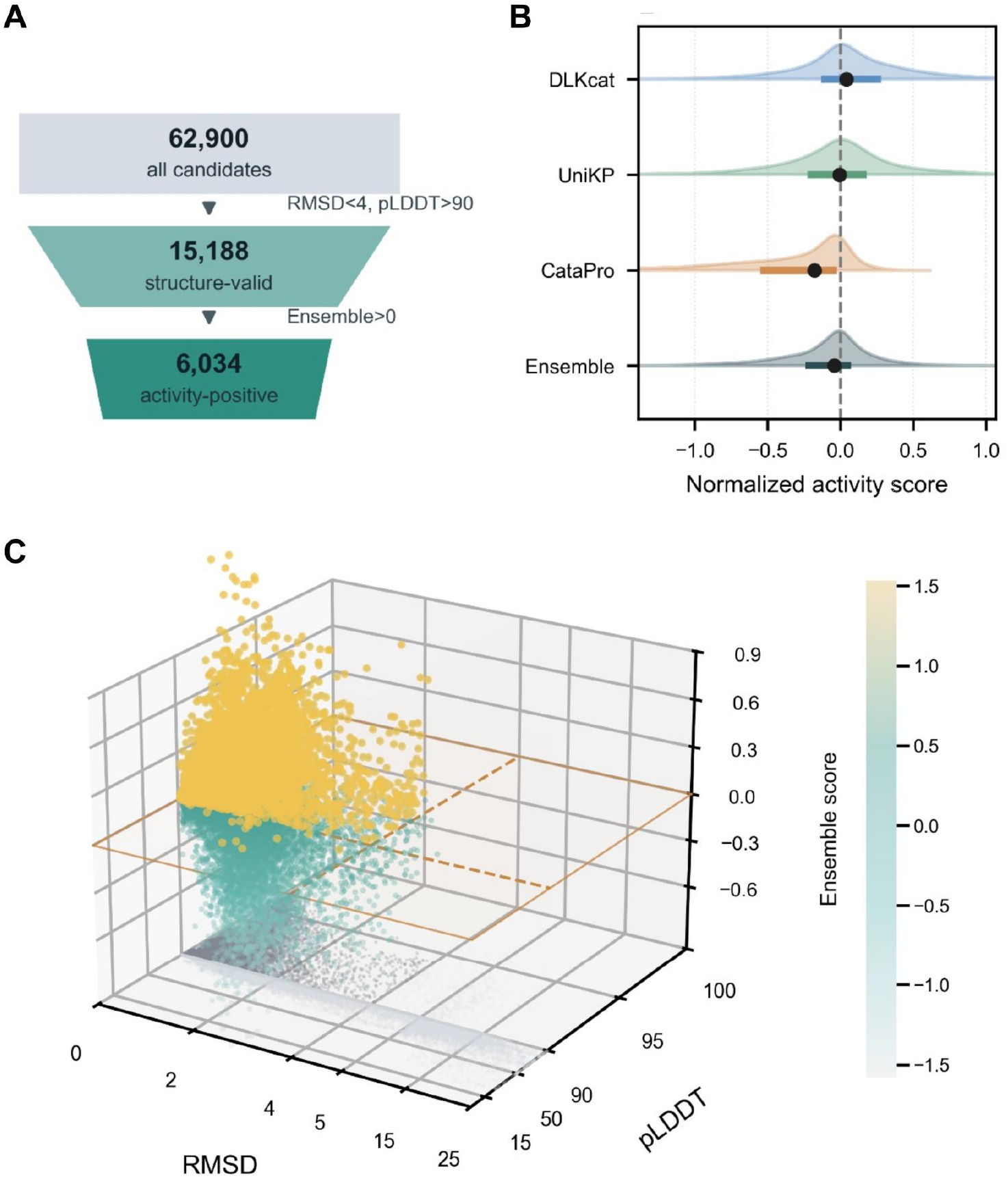
Generative Dataset Curation enriches structurally plausible and activity-positive variants. (A) Candidate filtering funnel. EnzymeIF generated 62,900 variants from 6,290 training-set backbones. ESMFold-based structural filtering retained 15,188 structure-valid variants satisfying backbone RMSD < 4 Å and pLDDT > 90. Ensemble activity filtering then retained 6,034 activity-positive variants with S_ensemble > 0. (B) Distributions of predictor-specific normalized activity scores for DLKcat, UniKP, and CataPro, together with the final ensemble score, computed over the structure-valid candidate pool. Dots indicate medians, thick horizontal bars indicate interquartile ranges, and thin horizontal bars indicate bootstrap confidence intervals of the median. (C) Structure-activity landscape of generated candidates in the backbone RMSD-pLDDT-ensemble score space. Candidates satisfying the structural gate are shown in darker gray, and activity-positive variants retained by GDC are highlighted. Threshold planes mark backbone RMSD = 4 Å, pLDDT = 90, and *S*_*ensemble*_ = 0.

The structural plausibility filter retained 15,188 variants, corresponding to 24.1% of the generated candidate pool, by requiring backbone RMSD < <4 Å and mean pLDDT > 90 after ESMFold refolding. These structure-valid variants were then scored by the DLKcat–UniKP–CataPro ensemble using predictor-normalized mutant–native activity differences. Applying the ensemble threshold *S*_ensemble_ > 0 yielded 6,034 activity-positive variants, corresponding to 9.6% of the original candidate pool. The marginal pLDDT and backbone-RMSD distributions for the full 62,890-variant candidate pool are shown in Figure S1B,C.

The resulting curated set defines an activity-enriched sequence distribution for supervised CatIF training. The funnel analysis in Figure 2A summarizes the reduction from all generated candidates to structure-valid and then activity-positive variants. Predictor-specific normalized score distributions and the final ensemble distribution are shown in Figure 2B. The structure–activity landscape in Figure 2C further indicates that activity-positive candidates occupy a constrained region jointly shaped by RMSD, pLDDT, and ensemble-predicted catalytic improvement. Together, these results show that GDC converts broad EnzymeIF sampling into a more selective training distribution enriched for variants that satisfy both structural plausibility and predicted activity-improvement criteria. (See also Table 1 for the upstream training-set partition this candidate pool was sampled from.)

### 3.3 CatIF-RL Achieves the Largest Predicted Catalytic Gain Among Inverse-Folding Baselines

We next benchmarked CatIF-RL against CatIF, EnzymeIF, the generic GraDe-IF reference^45^, and representative inverse-folding baselines, including ProteinMPNN^26^, LigandMPNN^28^, ESM-IF^48^, ABACUS-T^29^, and PiFold^27^. All models were evaluated on the DLKcat-aligned held-out benchmark using the unified paired mutant–native protocol described in Methods. The full benchmark summary is given in Table 2, with detailed per-protein statistics (mean ± 95 % bootstrap CI and paired Wilcoxon p-values) in Table S7.

CatIF-RL achieved the strongest shift toward improved predicted catalytic turnover. The final CatIF-RL checkpoint, corresponding to RL Round 3, reached a Δlog_10_ *k*_cat_ of 0.6049, exceeding CatIF (0.3647), EnzymeIF (0.2359), and GraDe-IF (0.1115). In contrast, several structure-oriented baselines yielded near-zero or negative predicted turnover shifts under the same paired evaluation protocol, including ESM-IF (0.0751), ABACUS-T (0.0061), LigandMPNN (-0.0550), PiFold (-0.0858), and ProteinMPNN (-0.1182). The per-protein distribution in Figure 3A shows the corresponding rightward displacement of CatIF-RL relative to the other models. Per-baseline paired-difference histograms (CatIF-RL vs each other model) are shown in Figure S8, and the recovery–activity scatter is shown in Figure S5.

**Figure 3.**
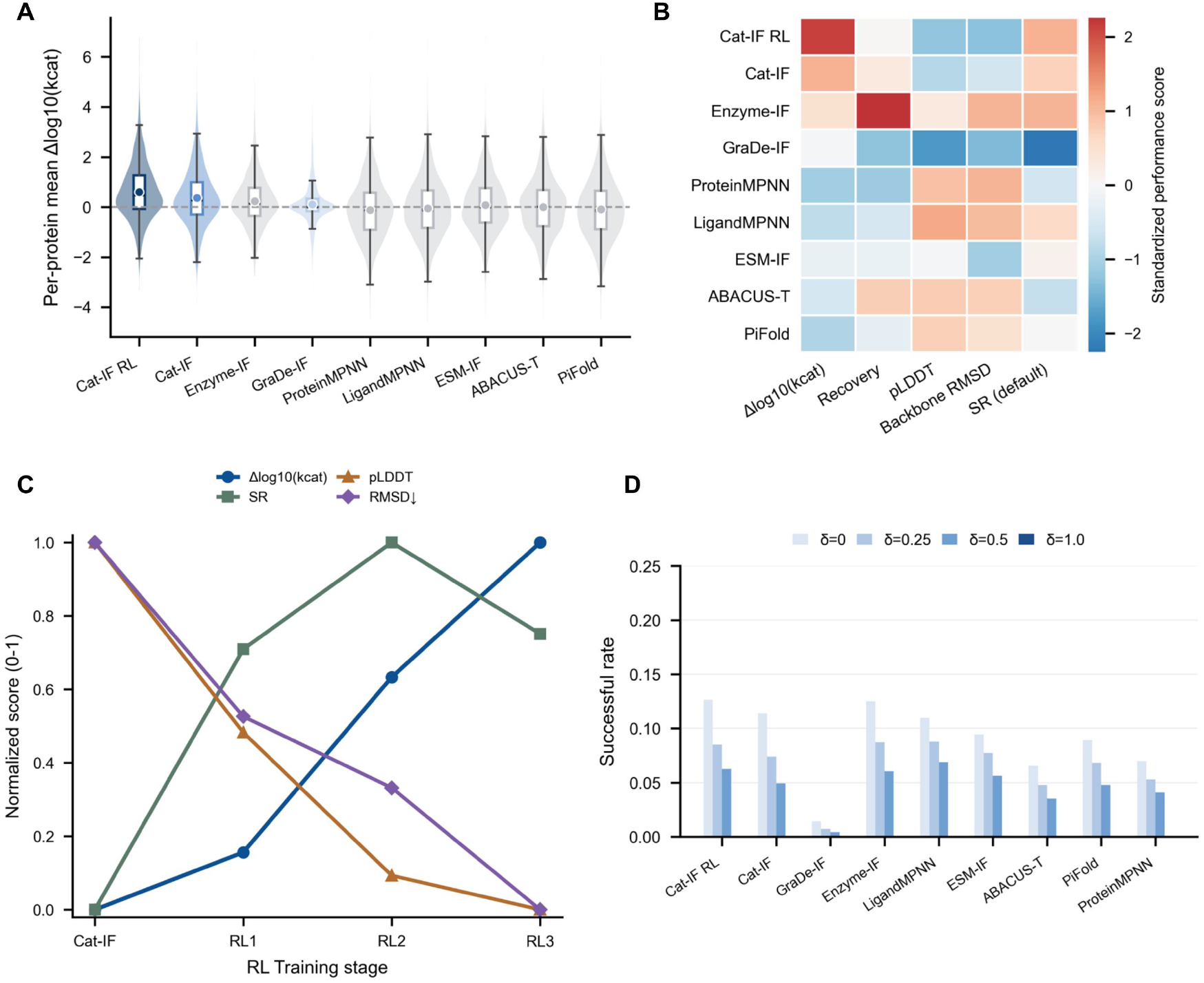
Benchmark comparison of CatIF-RL and representative inverse-folding baselines on the DLKcat-aligned held-out test set. (A) Distribution of per-protein predicted turnover improvement, measured as Δlog _10_ (*k*_cat_), across CatIF-RL, CatIF, EnzymeIF, GraDe-IF, and representative inverse-folding baselines. Violin plots show protein-level Δlog _10_ (*k*_cat_) distributions under paired mutant-native DLKcat evaluation. Boxplots indicate interquartile ranges and medians, and overlaid markers summarize mean values and confidence intervals where shown. (B) Standardized heatmap summarizing multi-metric benchmark performance across models. Displayed values are z-score-normalized across models, with backbone RMSD sign-inverted so that higher values uniformly indicate better performance. The heatmap includes Δlog _10_ (*k*_cat_), Recovery Rate, pLDDT, backbone RMSD, and default SR. (C) Stage-wise trajectory from CatIF through CatIF-RL Rounds 1 to 3. Curves summarize normalized trends for Δlog _10_ (*k*_cat_), default SR, pLDDT, and backbone RMSD, with RMSD converted to a higher-is-better orientation for visualization. (D) Success-rate comparison under increasingly stringent catalytic improvement thresholds. SR quantifies the fraction of generated variants simultaneously satisfying the structural plausibility criteria and the corresponding minimum Δlog _10_ (*k*_cat_) requirement.

The performance profile of CatIF-RL is more nuanced when structural and sequence-conservation metrics are considered jointly. CatIF-RL achieved a default SR@*δ*=0 of 0.1266, higher than CatIF (0.1145) and far above GraDe-IF (0.0141), indicating that activity-oriented post-training increases the fraction of variants satisfying the default joint structure–function criterion (Figure 3D). Joint success-rate sensitivity to *δ* ∈ {0, 0.25, 0.5, 1.0, 1.5, 2.0} is reported in Table S9. However, CatIF-RL did not dominate conservative design metrics. Its Recovery Rate was 0.5533, compared with 0.5719 for CatIF and 0.7088 for EnzymeIF. Its mean pLDDT was 77.87 and backbone RMSD was 5.3224 Å, whereas structure-oriented baselines such as LigandMPNN, ProteinMPNN, and ABACUS-T achieved higher pLDDT values or lower RMSD values. The compact heatmap in Figure 3B summarizes this multi-metric profile: CatIF-RL is strongest along the predicted activity axis, while more conservative inverse-folding baselines remain competitive in structural plausibility and sequence-conservation metrics.

**Significance of these multi-metric differences vs CatIF-RL is summarized in Figure S7, and the EC-class-stratified breakdown is shown in Figure S6**.

### 3.4 RL Rounds Reveal a Structure–Function Trade-Off

The round-wise trajectory from CatIF to CatIF-RL further clarifies how RL reshapes the sequence distribution. Starting from CatIF, which achieved Δlog _10_ *k*_cat_ = 0.3647, successive RL rounds increased predicted catalytic improvement to 0.4063 in Round 1, 0.5186 in Round 2, and 0.6049 in Round 3. Thus, the primary activity metric increased monotonically across RL rounds (Figure 3C). The corresponding inner-loop GRPO convergence curves (loss / KL proxy / icorr) for each round are shown in Figure S3.

This increase in predicted activity was accompanied by a gradual shift away from conservative sequence and structure metrics. Recovery Rate decreased from 0.5719 for CatIF to 0.5612, 0.5574, and 0.5533 across RL Rounds 1–3. Similarly, pLDDT decreased from 79.16 to 78.49, 77.99, and 77.87, while backbone RMSD increased from 4.8850 Å to 5.0923 Å, 5.1772 Å, and 5.3224 Å. In contrast, the default success rate changed only modestly across the same trajectory: SR@*δ*=0 was 0.1145 for CatIF and 0.1259, 0.1308, and 0.1266 for RL Rounds 1 to 3, respectively. (Figure 3C,D). Round-wise complete metrics — including ΔlgKcat, Recovery, pLDDT, backbone RMSD, and joint success rate at four *δ* thresholds with 95 % bootstrap CIs — are listed in Table S8, Figure 3.

These trends indicate that additional RL rounds primarily increase the average predicted activity of the generated candidate pool, rather than proportionally expanding the subset of variants that simultaneously satisfy stringent structural criteria. Thus, Δlog _10_ *k*_cat_ and threshold-based SR capture distinct aspects of design performance: the former measures the distributional shift toward higher predicted activity, whereas the latter measures the fraction of variants passing a joint function–structure decision rule.

### 3.5 Case Studies: Global Redesign and Motif-Preserving Inpainting

To complement the benchmark-level analysis, we examined three representative held-out global redesign cases and one motif-preserving local redesign case (Figure 4). These examples were evaluated using the same computational pipeline as the benchmark.

**Figure 4.**
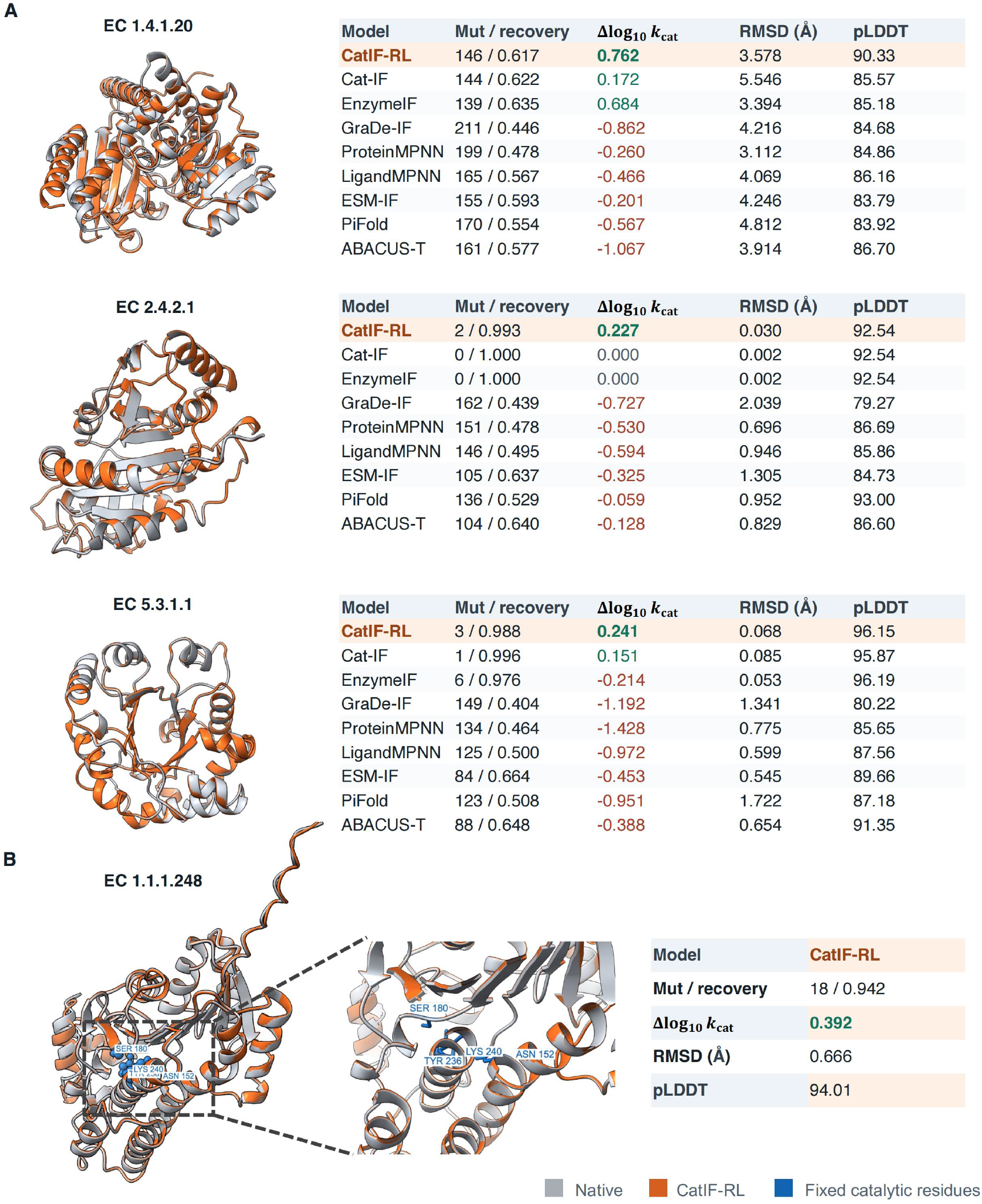
Representative case studies of CatIF-RL global redesign and motif-preserving inpainting. (A) Held-out EC entries redesigned under the unconstrained global protocol. Native-sequence reference structures are colored gray, and ESMFold-refolded CatIF-RL designs are colored orange. Tables summarize CatIF-RL, intermediate pipeline variants, and external inverse-folding baselines in terms of mutation count/recovery, Δlog _10_ (*k*_cat_), backbone RMSD, and pLDDT. EC 1.4.1.20 illustrates sequence-divergent redesign, whereas EC 2.4.2.1 and EC 5.3.1.1 illustrate near-native local editing. (B) Motif-preserving redesign of salutaridine reductase (EC 1.1.1.248) by subgraph inpainting. Catalytic residues (blue) were fixed based on prior mutagenesis evidence implicating these residues in the SalR catalytic/proton-transfer system, while surrounding positions were redesigned by CatIF-RL.

For the three global redesign cases, sequences were generated using the same unconstrained CatIF-RL sampling procedure, without target-specific steering, masking, or changes to the optimization objective. The different substitution counts therefore reflect target-dependent outputs of the same model rather than distinct design strategies. For EC 1.4.1.20, CatIF-RL introduced 146 substitutions, corresponding to a recovery of 0.617, and achieved Δlog _10_ *k*_cat_ = 0.762, the highest predicted activity gain among the displayed methods. EnzymeIF also showed a strong positive shift for this target (0.684), whereas CatIF reached 0.172, suggesting that this entry is relatively permissive to structure-conditioned redesign.

The other two global cases showed much smaller sequence deviations under the same protocol. For EC 2.4.2.1, CatIF-RL introduced two substitutions and achieved Δlog _10_ *k*_cat_ = 0.227, with recovery of 0.993, RMSD of 0.030 Å, and pLDDT of 92.54. For EC 5.3.1.1,CatIF-RL introduced three substitutions and achieved Δlog _10_ *k*_cat_ = 0.241, with recovery of 0.988, RMSD of 0.068 Å, and pLDDT of 96.15. These examples indicate that positive predicted activity shifts can arise with different extents of sequence change across targets. Per-baseline metrics for the four cases (mutation count, recovery, ΔlgKcat, RMSD, pLDDT) are listed in Table S11, and the same matrix is visualized in Figure S9.

We further evaluated motif-preserving local redesign using salutaridine reductase (EC 1.1.1.248). In this constrained setting, Asn152, Ser180, Tyr236, and Lys240 were fixed based on prior mutagenesis evidence implicating them in the SalR catalytic/proton-transfer system^49^, while surrounding positions were redesigned by subgraph inpainting. The four fixed motif residues with literature provenance are listed in Table S12. The resulting CatIF-RL design introduced 18 substitutions outside the preserved motif and retained a positive predicted activity shift, with Δlog _10_ *k*_cat_ = 0.392, recovery of 0.942, RMSD of 0.666 Å, and pLDDT of 94.01. Full case-level annotations and per-baseline outputs are provided in Table S11, and visualized in Figure S9.

## 4. Discussion

In this study, we reformulated a structure-conditioned discrete-diffusion inverse-folding framework into a generative enzyme-sequence design pipeline for improving catalytic activity. Using GraDe-IF, a graph-denoising diffusion model for inverse protein folding, as a generic architectural reference, we introduced three sequential components^45^. First, we adapted the inverse-folding prior to the enzyme domain through enzyme-focused training and structural regularization, yielding the enzyme-specialized model EnzymeIF. Second, we applied GDC to construct an activity-enriched variant set and trained CatIF from scratch on this curated distribution. Third, we further refined the CatIF sampling policy with KL-regularized GRPO, resulting in CatIF-RL^46^. In addition, we developed a motif-preserving partial-design variant based on a discrete subgraph inpainting sampler, enabling localized sequence redesign under explicit functional constraints^19,47,50,51^.

On a leakage-controlled DLKcat holdout benchmark evaluated under a unified protocol, CatIF-RL achieved a Δlog _10_ (*k*_cat_) of 0.6049, corresponding to an average predicted turnover more than four-fold higher than that of the corresponding native enzymes^13^. This value was substantially higher than its immediate no-RL counterpart CatIF (0.3647), the enzyme-specialized structure-only variant EnzymeIF (0.2359), and the generic GraDe-IF architectural reference (0.1115)^45^. The progression from GraDe-IF to the successive pipeline variants was monotonic (GraDe-IF → EnzymeIF → CatIF → CatIF-RL: 0.1115 → 0.2359 → 0.3647 → 0.6049). Within the controlled pipeline, the consecutive improvements from EnzymeIF to CatIF and then to CatIF-RL indicate that enzyme-domain adaptation, activity-enriched supervised training, and RL-based refinement contribute non-redundant effects. CatIF-RL also outperformed the other external inverse-folding baselines, several of which yielded negative predicted shifts^25–29^. Case-level analyses further showed that the same global sampling procedure can yield DLKcat-predicted gains with different degrees of sequence divergence across targets, while motif-preserving inpainting provides a constrained setting for redesigning sequence context around fixed catalytic residues^24,35,36^. The corresponding pairwise paired-Wilcoxon tests across all four metrics (each transition independently significant; BH-adjusted q < 0.001 for ΔlgKcat and Recovery) are reported in Table S10.

The performance of CatIF-RL is attributable to three sequential design choices, rather than to the simple application of RL to a generic inverse-folding model. A fourth component, the motif-preserving sampler, addresses the separate but practically important requirement of constrained enzyme redesign. The first design choice concerns enzyme-domain adaptation of the structural prior. The original GraDe-IF model treats general-purpose PDB structures as graph inputs without distinguishing enzymes from non-enzymes^45^. This setting is appropriate for general inverse folding, but is not specifically aligned with enzyme design, where catalytic function imposes additional constraints on the relevant sequence–structure distribution. To construct EnzymeIF, we used BRENDA-derived predicted enzyme structures as enzyme-domain examples and incorporated CATH-derived general protein structures during fine-tuning as structural regularization^44,52^. These non-enzymatic structures serve a dual role: they provide noise for denoising regularization and broaden geometric coverage of the training distribution. As a result, EnzymeIF acquired stronger enzyme-specialized inverse-folding capacity and improved tolerance to predicted structural inputs, as reflected by the increase in predicted mean Δlog _10_ (*k*_cat_) from 0.1115 for GraDe-IF to 0.2359 for EnzymeIF. The second design choice is activity-enriched supervised training through GDC. Rather than initializing RL directly from a structure-conditioned model, we first used an iterative generate–filter– retrain procedure to construct a variant set enriched for high predicted catalytic activity. Candidate variants were filtered using an ensemble of independent *k*_cat_ predictors together with structural-plausibility criteria^13–15,48^. CatIF was then trained from scratch on this curated activity-positive dataset. Consequently, before any RL signal was applied, the CatIF policy already assigned appreciable probability mass to regions of sequence space with higher predicted activity, raising Δlog _10_ (*k*_cat_) to 0.3647. RL then amplified this pre-enriched distribution rather than replacing the GDC training stage itself, further increasing Δlog _10_ (*k*_cat_) from 0.3647 to 0.6049. This staged design likely accounts for the improved sample efficiency of CatIF-RL relative to a direct RL procedure applied to a generic protein generator^37–39,43,46,53^. The consecutive gains from EnzymeIF, CatIF, and CatIF-RL therefore correspond to distinct levels of distributional reweighting: enzyme-domain adaptation, activity-enriched supervised reshaping, and policy-level refinement. The third design choice concerns the construction of the evaluation and reward signals. Because DLKcat contributes to both reward modeling and final evaluation, we strictly retained the original DLKcat train, development, and test partition to avoid circular assessment^13^. At the same time, the activity reward used during training was not derived from DLKcat alone, but from a scale-normalized ensemble of DLKcat, UniKP, and CataPro^13–15^. All activity estimates were expressed as mutant-minus-wild-type differences computed within the same predictor. Per-predictor normalization prevents any single model from dominating the reward solely because of its output scale, whereas the differential formulation controls for systematic calibration offsets within each predictor and partially decouples the design signal from any one predictor’s inductive bias. These choices do not eliminate surrogate-model uncertainty, but they reduce the susceptibility of the reported improvements to leakage, scale effects, or predictor-specific artifacts. For constrained design, we further introduced a discrete subgraph inpainting sampler. Fixed positions are sampled directly from the forward marginal conditioned on the ground-truth sequence, followed by iterative resampling to restore local compatibility near the boundary between constrained and designable regions. This design prevents fixed functional regions from inheriting unnecessary uncertainty from noisy intermediate states, making it better suited to local sequence redesign under fixed catalytic residues or other critical motifs. Conceptually similar inpainting strategies have been widely adopted in constrained generation, including general diffusion inpainting, motif scaffolding in protein structure generation, and CDR-focused antibody design^19,47,50,51^. Our implementation represents a discrete-sequence analogue of this methodological direction within a graph-based diffusion framework.

Beyond the overall increase in predicted catalytic activity, the results reveal a non-trivial trade-off between functional optimization and structural conservatism. CatIF-RL achieved the largest predicted mean functional gain but did not dominate conservative design metrics. Its sequence recovery was 0.5533, lower than the 0.7088 achieved by EnzymeIF; its pLDDT was 77.87, lower than 83.74 for EnzymeIF and 87.05 for LigandMPNN; and its backbone RMSD was 5.32 Å, higher than the 3.86 Å obtained by ProteinMPNN^26,28,48^. Similarly, under the strictest joint pass criterion, SR@*δ*=1.0, CatIF-RL achieved 0.0345, essentially comparable to LigandMPNN at 0.0338, despite the larger predicted activity shift of CatIF-RL. Importantly, this trade-off was not introduced solely by the RL stage. Already at the CatIF stage, Recovery Rate had decreased to 0.5719 and pLDDT to 79.16, values close to those observed for CatIF-RL. A more plausible interpretation is therefore that activity-oriented reweighting of the sequence distribution begins during GDC, whereas RL further amplifies this trend without fundamentally reshaping the design distribution. Round-wise behavior supports this interpretation: predicted mean Δlog _10_ (*k*_cat_) increased consistently from Round 1 to Round 3, whereas SR@*δ*=0 remained near 0.126 across all three rounds. Additional RL rounds therefore continued to shift the candidate pool toward higher predicted activity but did not proportionally expand the subset of candidates satisfying strict structural criteria.

These contrasting outcomes between average predicted improvement and threshold-based success rate point to a deeper distinction between the two metrics. The former measures the distributional shift of the generated candidate pool relative to the native enzyme in the direction of higher predicted activity, whereas the latter measures the fraction of candidates that pass a stringent joint functional–structural decision rule. This distinction matters in practice because pool-level optimization and single-candidate success correspond to different deployment modes: the former is suited to downstream ranking and screening, whereas the latter more closely approximates direct-hit design. Our results suggest that the region of sequence space satisfying both strong functional constraints and strong structural conservatism is relatively narrow. By contrast, when predicted activity alone is optimized, substantial improvements remain accessible across a broader sequence region. This observation suggests that the pipeline may be steering the policy toward sequence regions that are sparsely populated by natural enzymes but remain structurally plausible under the surrogate ensemble, consistent with guided exploration of dark sequence space^20,21^.

Several limitations qualify these conclusions. First, the entire pipeline relies on surrogate models for structural and activity assessment, including ESMFold for structural evaluation and DLKcat, UniKP, and CataPro for *k*_cat_ estimation^13–15,48^. Because non-negligible disagreement remains among predictors, the term “predicted improvement” should be interpreted as conditional on the current surrogate models of catalytic function. Second, the RL stage is not an end-to-end optimization through the full diffusion trajectory; it is closer to policy optimization over sampled sequences under a discrete-diffusion prior. This choice was made to improve training stability and sample efficiency, rather than to provide a theoretically exhaustive formulation of diffusion-model RL^37,43,46^. Third, the unified benchmark uses backbone-only inputs. Baselines such as LigandMPNN or ABACUS-T may perform more strongly when supplied with richer task-specific inputs, such as ligand context, cofactors, multiple conformations, or MSA information^28,29^. Accordingly, our conclusions should be interpreted within this benchmark regime. Finally, all reported improvements are computational. Experimental validation of selected candidates remains necessary to establish the practical design capacity of the model. The pairwise predictor agreement (Pearson r ranges 0.13–0.59 over 25,970 matched triplets) is shown in Figure S4.

In conclusion, CatIF-RL illustrates a function-aware protein design paradigm built on an inverse-folding prior. Its central contribution is not the addition of reinforcement learning at the end of a generative model, but a staged procedure that combines enzyme-domain adaptation, activity-enriched supervised training through GDC, and KL-regularized policy optimization under a structure-conditioned discrete-diffusion prior^24,35– 39,43,45,46,53^. Through this procedure, an inverse-folding distribution originally oriented toward native-sequence recovery can be systematically reweighted toward regions defined by catalytic function. The resulting pipeline preserves the controllability of structure-conditioned generation while introducing an explicit functional bias through data curation and reward optimization. In this sense, inverse folding is extended from the recovery of plausible sequences for a given backbone to the search for functionally improved sequences under structural constraints. This paradigm is also extensible. In principle, the same framework can be generalized from single-objective *k*_cat_ optimization to multi-objective or uncertainty-aware rewards, for example by incorporating RMSD, pLDDT, or predictor disagreement into the RL objective together with *k*_cat_^10,12-14,48^. It can also be adapted from catalytic turnover to other design targets, including thermostability, substrate specificity, solubility, and expression feasibility^10^. As structural and functional surrogate models continue to improve, more accurate predictors can be substituted into the same filtering and reward interfaces without changing the overall training scheme^10,12-14,48^. CatIF-RL provides not only a concrete model for catalytic-activity-oriented enzyme design, but also a generalizable template for guiding protein inverse folding toward definable, composable, and constraint-aware functional objectives. (See Algorithm S1, Algorithm S2, and the SI Methods §S1 for full procedural detail.)

## ASSOCIATED CONTENT

### Supporting Information

The Supporting Information is available. CatIF-RL_SI.pdf — Extended methods (Section S1), training hyperparameters (Tables S1–S3), dataset and split statistics (Table S4), predictor ensemble details (Table S5), per-baseline sampling configuration (Table S6), pseudocode for the BLOSUM iterative-fusion variant (Algorithm S1) and the GRPO update (Algorithm S2), and supporting figures (Figures S1–S9).

### Data Availability

The CatIF-RL implementation, training scripts, data preparation pipeline, GRPO update routine, pre-trained model weights, and processed datasets are available at https://github.com/lynk-101-li/CatIF-RL.

## AUTHOR INFORMATION

### Authors

Yanheng Li, Key Laboratory of Molecular Medicine and Biotherapy in the Ministry of Industry and Information Technology, Department of Neurobiology, School of Life Sciences, Beijing Institute of Technology, Beijing 100081, China

Jialong Xiong, Key Laboratory of Molecular Medicine and Biotherapy in the Ministry of Industry and Information Technology, Department of Neurobiology, School of Life Sciences, Beijing Institute of Technology, Beijing 100081, China

Yuxin Zhang, Key Laboratory of Molecular Medicine and Biotherapy in the Ministry of Industry and Information Technology, Department of Neurobiology, School of Life Sciences, Beijing Institute of Technology, Beijing 100081, China

Tong Cai, Key Laboratory of Molecular Medicine and Biotherapy in the Ministry of Industry and Information Technology, Department of Neurobiology, School of Life Sciences, Beijing Institute of Technology, Beijing 100081, China

Chuan Fu, Key Laboratory of Molecular Medicine and Biotherapy in the Ministry of Industry and Information Technology, Department of Neurobiology, School of Life Sciences, Beijing Institute of Technology, Beijing 100081, China; Department of Orthopaedic Surgery, The Second Hospital of Shanxi Medical University, 382 Wuyi Road, Taiyuan, Shanxi 030001, China

Shutong Li, Key Laboratory of Molecular Medicine and Biotherapy in the Ministry of Industry and Information Technology, Department of Neurobiology, School of Life Sciences, Beijing Institute of Technology, Beijing 100081, China

Wei Xu, Key Laboratory of Molecular Medicine and Biotherapy in the Ministry of Industry and Information Technology, Department of Neurobiology, School of Life Sciences, Beijing Institute of Technology, Beijing 100081, China

Ruoyi Lyu, Key Laboratory of Molecular Medicine and Biotherapy in the Ministry of Industry and Information Technology, Department of Neurobiology, School of Life Sciences, Beijing Institute of Technology, Beijing 100081, China

Zhaoyang Chen, Key Laboratory of Molecular Medicine and Biotherapy in the Ministry of Industry and Information Technology, Department of Neurobiology, School of Life Sciences, Beijing Institute of Technology, Beijing 100081, China

Zheng Guo, Key Laboratory of Molecular Medicine and Biotherapy in the Ministry of Industry and Information Technology, Department of Neurobiology, School of Life Sciences, Beijing Institute of Technology, Beijing 100081, China

## ACKNOWLEDGMENTS

This work was supported by the National Natural Science Foundation of China (General Program), Grant No. 32571511 (to Feng Wang); the National Natural Science Foundation of China, Grant No. 82302705 (to Chuan Fu); and the Doctoral Fund Project of the Second Hospital of Shanxi Medical University, Grant No. 202301-13 (to Chuan Fu). The authors declare no competing financial interest. The authors used OpenAI ChatGPT to assist with language polishing of selected manuscript sections. All scientific content, claims, and interpretations are the sole responsibility of the authors, who reviewed and edited all AI-assisted text.

## Notes

https://github.com/lynk-101-li/CatIF-RL

### Competing Interest Statement

The authors have declared no competing interest.

### Summary of Updates

Changes from version 1: 1. Author list expanded from 6 to 12 authors. Six additional co-authors (Chuan Fu, Shutong Li, Wei Xu, Ruoyi Lyu, Zhaoyang Chen, Zheng Guo) contributed to quality-control verification, code repository organization and reproducibility testing, and manuscript review. This update aligns the preprint author list with the manuscript currently under review at J. Chem. Inf. Model. (ACS). 2. A third affiliation has been added (Department of Orthopaedic Surgery, The Second Hospital of Shanxi Medical University) for Chuan Fu's clinical affiliation. 3. Funding information updated: in addition to NSFC Grant 32571511 (Feng Wang), two grants supporting Chuan Fu's contribution have been disclosed - NSFC Grant 82302705 and the Doctoral Fund Project of the Second Hospital of Shanxi Medical University (Grant 202301-13). 4. Minor wording fixes throughout (no changes to scientific results, methods, figures, tables, or conclusions of the main text). 5. Supporting Information PDF: table styling refreshed for improved readability; content unchanged (all Tables S1~S6, Figures S1~S9, and Algorithms S1~S2 retained). All co-authors have consented to this revision and to the JCIM submission.

