## Supporting Information for "CatIF-RL: Activity-Oriented Enzyme Sequence Design by Steered Inverse Protein Folding"

### S1 Supplementary Methods

#### S1.1 Dataset construction details

The DLKcat–BRENDA enzyme-kinetic dataset (Li et al. [12]; BRENDA database, Chang et al. [40]) and the leakage-controlled train / validation / test split summarised in manuscript §2.1 are documented in full in **Table S4**. Sequence-level filtering proceeded in the manuscript-stated order — length cap (> 1,180 AA), removal of non-standard residues, and de-duplication of identical protein sequences — collapsing 16,839 BRENDA records to 7,713 distinct enzyme sequences (Table S4(a)). **Figure S1A** visualises the resulting per-sequence length distribution, including the 1,180-AA cutoff. CATH v4.2.0 backbones (Sillitoe et al. [47]), used **only** as structural regularisers during EnzymeIF training (manuscript §2.2), are kept entirely separate from the held-out 1,423-enzyme benchmark. All ESMFold-predicted backbones (Lin et al. [48]) used as graph inputs were generated with a fixed random seed (1234) splitting the enzyme set 9:1 between train and validation.

#### S1.2 EnzymeIF training details

EnzymeIF (manuscript §2.2) was trained on 23,682 backbones (5,661 enzymes + 18,021 CATH structures) under the GraDe-IF discrete-diffusion architecture (Yi et al. [41]). **Table S1** lists the full hyperparameter set. Key operational choices are: 6 EGNN layers, hidden dimension 128, 500 diffusion timesteps with a BLOSUM transition kernel, Adam learning rate $5 \times 10^{-4}$, weight decay $1 \times 10^{-5}$, batch size 32, gradient clipping at maximum norm 1.0, and exponential-moving-average decay 0.995. The selected checkpoint (epoch 467) corresponds to the lowest validation cross-entropy loss; **Figure S2A** shows the corresponding training and validation loss together with validation recovery rate and scaled perplexity.

#### S1.3 CatIF training details

CatIF (manuscript §2.4) was trained **from scratch** on the 6,034 activity- positive GDC variants — it is not a continuation of, nor distillation from EnzymeIF. CatIF reuses the EnzymeIF architecture and optimiser exactly; the only differences are the training distribution and the run identifier.

**Table S2** lists the full hyperparameter set; **Figure S2B** shows the resulting training and validation curves, with the round-end checkpoint selected at epoch 228 (lowest validation loss).

#### S1.4 GRPO training details

The KL-regularised GRPO refinement of CatIF (manuscript §2.5; GRPO objective introduced in Guo et al. [42]) is fully specified in **Algorithm S2** as a two-level loop: an outer round loop ($K = 3$ rounds of sample → score → train, each round generating fresh offline samples for the inner loop) and an inner GRPO update loop that operates on pre-scored CSVs without invoking the predictor ensemble during back-propagation. Notable operational choices that are flagged in the inner-loop pseudocode include (i) within-group sequence deduplication and a minimum of three distinct rewards per group; (ii) length-normalised token-mean log-prob to avoid length bias; (iii) a clipped MSE proxy for the KL term, with a multiplicatively adapted β bounded in [$\beta_{min} = 5 \times 10^{-4}$, $\beta_{max} = 0.5$] (iv) a mutation-budget penalty $\lambda_{\text{mut}} \cdot \max(0, \mu - \mu_0)$ activated only above the free-mutation threshold $\mu_0 = 0.30$.

**Table S3** lists the full hyperparameter set across all three rounds — they are identical except for the inner-epoch budget (the epoch-02 checkpoint of every round was selected for downstream use).

**Figure S3** shows total loss, KL proxy, and the reward × log-prob correlation (icorr) as a function of inner epoch for each round.

#### S1.5 Subgraph inpainting algorithm

Motif-preserving partial sequence design (manuscript §2.6) is implemented as RePaint-style (Lugmayr et al. [43]) discrete subgraph inpainting on top of the CatIF (or CatIF-RL) discrete-diffusion policy. **Algorithm S1** gives the exact procedure: for every reverse-step ($t \rightarrow s$) the wild-type sequence is forward-corrupted to step $s$ and fused with the model-predicted reverse step under a residue-level mask $m$, with $m_i = 1$ fixing position $i$ to wild-type and $m_i = 0$ marking it designable. Because CatIF uses a BLOSUM transition kernel, the closed-form jump-back operation of standard RePaint (uniform-noise variant) is not available; the case- study runs therefore use the iterative-fusion variant ($U = 5$) in which the denoise → GT-injection step is repeated $U$ times against the unchanged mask at every outer step, progressively refining boundary compatibility.

#### S1.6 Baseline reproduction details

The five external inverse-folding baselines (ProteinMPNN [25], ESM-IF [24], LigandMPNN [27], PiFold [26], ABACUS-T [28]) and the four CatIF-pipeline variants (GraDe-IF [41], EnzymeIF, CatIF, CatIF-RL) were sampled on the same 1,423-enzyme test set under the protocol summarised in manuscript §2.7.

**Table S6** lists the per-baseline sampling configuration, with seeds standardised to {1111, 2222, 3333, 4444, 5555} (5 seeds × 1 design per seed = 5 designs per backbone for every method). **Table S5** records the version, repository, conda environment, and inference interface of the three k_cat predictors (DLKcat [12], UniKP [13], CataPro [14]) used both for activity reward (manuscript §2.5) and held-out evaluation (manuscript §2.7). As stated in manuscript §2.7, structural metrics (pLDDT, backbone RMSD, joint success rate) were computed on a single-seed pre-specified ESMFold- refolded set [48] rather than on all five sampling rounds, to keep refolding compute tractable. All inference, GDC scoring, CatIF supervised training, and CatIF-RL refinement reported here were carried out on a single local NVIDIA RTX 5060 Ti 16 GB GPU; only EnzymeIF supervised training was performed on an NVIDIA RTX 4090 24 GB cloud GPU (AutoDL).

#### S1.7 Evaluation and statistics

The held-out benchmark statistics reported in manuscript Table 2 and §3.3 are reproduced and extended in **Table S7** (mean ± 95% bootstrap CI for all four metrics across all eleven baselines, with paired Wilcoxon p / BH-FDR- adjusted q vs CatIF-RL R3). Round-wise behaviour underlying §3.4 is captured in **Table S8** (CatIF / R1 / R2 / R3 across all four metrics plus joint success rate at four δ thresholds). **Table S9** reports the threshold-sensitivity analysis (SR @ $\delta \in \{0, 0.25, 0.5, 1.0, 1.5, 2.0\}$).

**Table S10** provides the ablation-pairwise significance test set (GraDe-IF → EnzymeIF → CatIF, plus R1 ↔ R2 ↔ R3) that supports the manuscript §4 claim of non-redundant pipeline contributions. **Figures S5, S7, and S8** visualise the same per-protein paired comparisons in three complementary forms: per-protein scatter (Recovery vs Δlog k_cat), multi-metric forest plot, and per-protein paired difference histograms.

**Figure S4** independently quantifies the inter-predictor agreement that motivates the ensemble formulation, and **Figure S6** stratifies the benchmark by EC main class to show that the pipeline trade-off is not class-specific.

#### S1.8 Case-study protocol

The four representative cases reported in manuscript §3.5 (three global redesigns and one motif-preserving SalR redesign) are detailed in **Table S11**. For consistency with the per-case values quoted in the manuscript narrative, Table S11 reports single-seed showcase values matching manuscript §3.5.

**Figure S9** visualises the full Table S11 metric matrix as a 4 × 4 grid of horizontal bar charts (rows = metric, columns = case), with CatIF-RL highlighted in every panel. **Table S12** lists the four fixed motif residues used for the SalR (EC 1.1.1.248) inpainting case (Asn152, Ser180, Tyr236, Lys240), per the catalytic / proton-transfer assignments of Geissler et al. [44]. Note that for the three global cases all baselines were sampled under the same unconstrained protocol; for SalR only CatIF-RL applied the motif constraint, while the other rows in Figure S9 column 4 are reference designs under the unconstrained protocol and therefore should not be compared on equal terms with the constrained CatIF-RL design.

### S2 Supplementary Tables

#### Table S1. EnzymeIF supervised training configuration.

EnzymeIF was trained on 23,682 enzyme + CATH structures (Table S4) with the GraDe-IF discrete diffusion architecture. Selected checkpoint (epoch 467 by training-script counter).

| Group | Hyperparameter | Symbol / flag | Value |
|---|---|---|---|
| *Architecture* | Backbone | — | EGNN denoiser (GraDe-IF) |
| | Number of EGNN layers | --depth | 6 |
| | Hidden channel dim | --hidden | 128 |
| | Residue embedding | --embedding | enabled |
| | Embedding dim | --embedding_dim | 128 |
| | Feature normalisation | --norm_feat | enabled |
| | Secondary-structure embed | --embed_ss (default) | -1 (disabled) |
| | Discrete transition kernel | --noise_type | BLOSUM |
| | Total diffusion timesteps | T | 500 |
| | Training objective | — | residue-level cross-entropy on predicted clean amino-acid distribution |
| *Optimiser* | Optimiser | — | Adam |
| | Learning rate | --lr | $5 \times 10^{-4}$ |
| | Weight decay | --wd | $1 \times 10^{-5}$ |
| | Dropout | --drop_out | 0.1 |
| | Batch size | --batch_size | 32 |
| | Gradient clipping (L2 max-norm) | — | 1.0 |
| | EMA decay | — | 0.995 |
| *Run* | Hardware | — | NVIDIA RTX 4090 24 GB (AutoDL cloud) |
| | Selected checkpoint epoch | — | 467 |
| | Checkpoint-selection criterion | — | lowest validation loss |

#### Table S2. CatIF supervised training configuration.

CatIF was trained **from scratch** on the 6,034 GDC-curated activity-positive variants (manuscript §2.3, §2.4) with the same architecture and optimiser as EnzymeIF. Selected checkpoint (epoch 228).

| Group | Hyperparameter | Symbol / flag | Value |
|---|---|---|---|
| *Architecture* | (Identical to Table S1) | — | — |
| | Number of EGNN layers | --depth | 6 |
| | Hidden channel dim | --hidden_dim | 128 |
| | Residue embedding | --embedding | enabled |
| | Embedding dim | --embedding_dim | 128 |
| | Feature normalisation | --norm_feat | enabled |
| | Secondary-structure embed | --embed_ss (default) | -1 (disabled) |
| | Discrete transition kernel | --noise_type | BLOSUM |
| | Total diffusion timesteps | T | 500 |
| *Optimiser* | Optimiser | — | Adam |
| | Learning rate | --lr | $5 \times 10^{-4}$ |
| | Weight decay | --wd | $1 \times 10^{-5}$ |
| | Dropout | --drop_out | 0.1 |
| | Batch size | --batch_size | 32 |
| | Gradient clipping (L2 max-norm) | — | 1.0 |
| | EMA decay | — | 0.995 |
| *Run* | Hardware | — | NVIDIA RTX 5060 Ti 16 GB (local) |

| | | |
|---|---|---|
| Selected checkpoint epoch | — | 228 |
| Initialisation | — | from-scratch (not warm-started from EnzymeIF) |

**Notes (apply to both tables)**

1. **EnzymeIF vs CatIF differ only in dataset, run-tag, and selected epoch.** All architecture and optimiser hyperparameters are identical between EnzymeIF and CatIF; the only actual differences are (i) the training distribution (Table S4) and (ii) which epoch's checkpoint was selected as the round-end model. 2. **Checkpoint selection.** Both EnzymeIF (epoch 467) and CatIF (epoch 228) were chosen as the epoch with the lowest validation loss. 3. **Sampling-time hyperparameters** (DDIM step, diverse-sampling flag, evaluation-time sampling step) are not in this table — they belong in Table S6 (baseline sampling configurations), where CatIF is one row.

#### Table S3. CatIF-RL (offline GRPO) training hyperparameters across the three iterative rounds.

All three rounds share an identical optimizer/loss configuration. The operational inner-epoch budget is **2 epochs per round** for all three rounds; the reference policy `π_ref` is the original CatIF checkpoint and is **frozen across all rounds**.

| Hyperparameter | Symbol | Round 1 | Round 2 | Round 3 |
|---|---|---|---|---|
| **Outer-loop sampling** | | | | |
| Group size per (enzyme, substrate) pair | G | 5 | 5 | 5 |
| DDIM step (sampling) | — | 100 | 100 | 100 |
| Sampling seed | — | 11 | 11 | 11 |
| Diverse decoding (categorical) | — | ON | ON | ON |
| GRPO inner loop — optimisation | | | | |
| Inner epochs per round (operational) | E_k | 2 | 2 | 2 |
| Training-run length (--epochs flag) | — | 50 | 25 | 25 |
| Warm-up epochs | — | 0 | 0 | 0 |
| Learning rate | $\eta$ | $5 \times 10^{-6}$ | $5 \times 10^{-6}$ | $5 \times 10^{-6}$ |
| Weight decay | wd | $1 \times 10^{-2}$ | $1 \times 10^{-2}$ | $1 \times 10^{-2}$ |
| AdamW betas | $(\beta_1, \beta_2)$ | (0.9, 0.98) | (0.9, 0.98) | (0.9, 0.98) |
| Gradient accumulation steps | accum | 1 | 1 | 1 |
| Gradient clipping (norm) | clip_g | 2.0 | 2.0 | 2.0 |
| AMP / fp16 | — | enabled | enabled | enabled |
| Random seed (training) | — | 42 | 42 | 42 |
| **GRPO inner loop — reward & KL** | | | | |
| Reward shaping mode | — | lin_sym | lin_sym | lin_sym |
| Reward scale | $\tau$ | 0.40 | 0.40 | 0.40 |
| Mutation penalty weight | λ_mut | 0.10 | 0.10 | 0.10 |
| Mutation-free fraction | $\mu_0$ | 0.30 | 0.30 | 0.30 |
| KL target | KL_target | 0.01 | 0.01 | 0.01 |
| KL clip (per-token logp gap) | c | 5.0 | 5.0 | 5.0 |
| Initial β | $\beta_0$ | 0.01 | 0.01 | 0.01 |
| β upper bound | β_max | 0.50 | 0.50 | 0.50 |
| β lower bound | β_min | $5 \times 10^{-4}$ | $5 \times 10^{-4}$ | $5 \times 10^{-4}$ |
| β multiplicative update (above target) | — | × 1.5 | × 1.5 | × 1.5 |
| β multiplicative update (below target) | — | × 0.7 | × 0.7 | × 0.7 |
| **GRPO inner loop — group filtering** | | | | |
| Min distinct rewards per group | R_min | 3 | 3 | 3 |
| Within-group sequence dedup | — | ON | ON | ON |
| Forward pass (logp evaluation) | | | | |
| Sampling timestep for logp | step | 100 | 100 | 100 |
| Checkpointing | | | | |
| π_φ initial checkpoint | — | $\pi_0$ (CatIF) | $\pi_1$ epoch 02 | $\pi_2$ epoch 02 |
| π_ref checkpoint (frozen) | — | $\pi_0$ (CatIF) | $\pi_0$ (CatIF) | $\pi_0$ (CatIF) |
| Save every | — | 1 epoch | 1 epoch | 1 epoch |
| Round-end checkpoint used downstream | — | epoch 02 | epoch 02 | epoch 02 |

**Notes.**

1. **Operational inner-epoch budget = 2.** Because GRPO is run offline against a fixed batch of pre-scored samples per round, training was capped at 2 epochs per round to mitigate over-fitting to that fixed batch. The training run was allowed to continue beyond 2 epochs only to inspect convergence behaviour in the diagnostic logs; the **epoch-02 checkpoint is consistently used as the round-end policy** that feeds the next

round and the test-set evaluations. The 2-epoch choice was conservative and was not benchmarked against longer training. 2. **Compute.** All CatIF-RL training was performed on a single NVIDIA RTX 5060 Ti GPU (16 GB VRAM). 3. **Same data set across rounds.** All three rounds source the conditioning graphs from the same BRENDA train + valid split; the difference between rounds is the generated sample CSV (round-$k$ samples are produced by $\pi_{k-1}$).

#### Table S4. Dataset construction and leakage-control details.

The independent benchmark of 1,423 enzymes was held out throughout the study. CATH v4.2.0 backbones served only as general-protein structural regularisers during EnzymeIF training (Section 2.2) and were not introduced into the held-out benchmark.

##### S4(a) Sequence-level filtering pipeline (DLKcat-BRENDA → 7,713 distinct enzymes)

Per-step removal counts: length > 1,180 AA removed 76 unique sequences; non-standard amino acids removed an additional 18 unique sequences (see Figure S1A for the corresponding length distribution).

| Step | Operation | Sequences retained | Δ |
|---|---|---|---|
| 0 | Initial DLKcat–BRENDA records (organism × substrate × sequence) | 16,839 | — |
| 1 | Remove sequences with length > 1,180 AA | 16,763 | −76 |
| 2 | Remove sequences containing non-standard residues | 16,745 | −18 |
| 3 | De-duplicate identical protein sequences | 7,713 | −9,032 |
| 4 | Predict ESMFold backbones (one per distinct sequence) | 7,713 backbone graphs | — |

##### S4(b) Train / validation / test split (preserving DLKcat split strategy[12])

| Train | Validation | Test (held-out benchmark) | Total |
|---|---|---|---|
| 5,661 | 629 | 1,423 | 7,713 |
| 18,021 | 608 | — | 18,629 |
| 23,682 | 1,237 | 1,423 | 26,342 |

##### S4(c) Leakage-control mechanics

| Mechanism | Implementation |
|---|---|
| Test-set lock | The 1,423 DLKcat-test sequences were assigned to the held-out benchmark before any model training. They are never used in EnzymeIF / CatIF / CatIF-RL training, GDC variant scoring, or RL reward computation. |
| Train/valid 9:1 split (enzymes only) | Within the 6,290 non-test enzymes, samples are split 5,661 / 629 by random shuffling with seed = 1234 and a 0.9 train ratio. |
| File-name uniqueness across {train, valid, test} | The dataset-construction pipeline skips any file whose name already appears in another split, guaranteeing disjoint train/validation/test sets. |
| Sequence-level test isolation | The DLKcat protein-cluster identifier (distc_pro_num) is propagated through the entire pipeline so that distinct-sequence variants of any test-set cluster cannot leak into train or validation. |
| CATH regularisers do not enter test | CATH backbones are added only into train and validation directories; the test directory contains 100 % DLKcat-test enzymes. |

##### S4(d) Data formats and preprocessing

| Item | Description |
|---|---|
| Per-protein graph file | .pt (PyTorch Geometric Data object) |
| Node features | one-hot residue identity (K = 20) + extra physicochemical/geometric features |
| Backbone source | ESMFold predicted structure for DLKcat–BRENDA enzymes; PDB-derived for CATH v4.2.0 |
| Random seed for splitting | 1234 |

#### Notes

1. **Source-of-truth consistency.** The four sub-tables together describe the dataset construction reported in manuscript §2.1 and Table 1. Numbers are taken verbatim from manuscript Table 1 and verified against the pipeline-asserted partition counts (6,290 → 5,661 / 629). 2. **CATH regularisation purpose.** As explained in §2.2, CATH structures broaden backbone-geometry coverage during EnzymeIF training but are never used to compute Δlog k_cat or related activity metrics, since they lack a substrate label. 3. **Reproducibility.** Re-running the full pipeline requires (i) the DLKcat-BRENDA raw records, (ii) the DLKcat test-protein

cluster identifier list (with the distc_pro_num field), (iii) ESMFold for backbone prediction, and (iv) the dataset-splitting and isolation pipeline (sequence-level filtering followed by 9:1 train/validation random shuffling). All scripts are available at the GitHub repository linked in the main manuscript's Code Availability statement.

#### Table S5. The three $k_{cat}$ predictors used to build $S_{ensemble}$.

For every (substrate × candidate sequence) pair, the three predictors `m ∈ {DLKcat, UniKP, CataPro}` independently produce $log_{10}k_{cat}$ . Their normalised Δ-scores are then averaged into `S_ensemble` (manuscript §2.3).

| Predictor | Citation | Original publication | Repository | Architecture (one-liner) | Input modalities | Output | Used at stage |
|---|---|---|---|---|---|---|---|
| DLKcat | Li et al., 2022 [12] | _Nat. Catal._ 5, 662–672 (2022). doi:10.1038/s41929-022-00798-z | https://github.com/SysBioChalmers/DLKcat | CNN over substrate + GNN over protein sequence | substrate SMILES + protein sequence | $log_{10}k_{ca}$ ($s^{-1}$) | GDC + RL reward + test eval |
| UniKP | Yu et al., 2023 [13] | _Nat. Commun._ 14, 8211 (2023). doi:10.1038/s41467-023-44113-1 | https://github.com/Luo-SynBioLab/UniKP | ProtT5 sequence embedding + SMILES Transformer + ExtraTrees regressor | substrate SMILES + protein sequence | $log_{10}k_{ca}$ | GDC + RL reward + test eval |
| CataPro | Wang et al., 2025 [14] | _Nat. Commun._ 16 (1), 2736 (2025). doi:10.1038/s41467-025-58038-4 | https://github.com/zchwang/CataPro | dual-branch deep-learning regressor combining a protein-language-model embedding with a chemistry encoder, trained for robust enzyme k_cat prediction | substrate SMILES + protein sequence | $log_{10}k_{ca}$ | GDC + RL reward + test eval |

##### Ensemble construction (manuscript §2.3)

For predictor `m` and (sequence x, substrate s):

$\Delta_m(x,s) = log_{10}\hat{k}_c at, m(x,s) - log_{10}\hat{k}_c at, m(x_n ative, s)\tilde{z}_m(x,s) = \Delta_m(x,s)/\left(q_{0.9}^{(m)} - q_{0.1}^{(m)}\right)$ (robust quantile-range scaling over the 15,188 structure-valid candidate pool) $S_{ensemble}(x,s) = (1/3) \cdot \Sigma_m \tilde{z}_m(x,s)$

$q_{0.9}^{(m)}$ and $q_{0.1}^{(m)}$ denote the 90th / 10th percentiles of $\Delta_m$ per predictor; they are estimated **once** on the structure-valid GDC candidate pool and reused for all downstream RL reward computations.

#### Notes

1. **All three predictors operate on (sequence, SMILES) — no structural input required.** This makes them cheap to call inside the GDC loop and the RL reward; structural plausibility is gated separately by ESMFold + RMSD/pLDDT thresholds (manuscript §2.3). 2. **Robust quantile-range normalisation.** The denominator $q_{0.9} - q_{0.1}$ is a robust scale estimator (less sensitive to outliers than σ); it does **not** subtract the median, so the sign of $S_{ensemble}$ is preserved relative to the native sequence. 3. **Ensemble freezing.** The three predictors and their quantile parameters are frozen at the start of GDC and kept identical through all CatIF supervised training, GRPO rounds 1–3, and final test-set evaluation. This avoids reward drift across rounds.

#### Table S6. Sampling configuration on the 1,423-enzyme independent benchmark — by baseline.

For all methods, sampling on the held-out benchmark used **5 independent random seeds: {1111, 2222, 3333, 4444, 5555}**, producing one designed sequence per (backbone, seed) and 5 sequences per backbone in total (7,115 sequences per method).

| Method | Reference | Backbone-conditioning input | Sampling protocol | Decoding parameters | Output format |
|---|---|---|---|---|---|
| ProteinMPNN | Dauparas et al., 2022[25] | PDB parsed to JSONL via the ProteinMPNN preprocessing helper | autoregressive masked decoding | --num_seq_per_target 1, --batch_size 1, --simple_output 1; default sampling temperature | FASTA |
| ESM-IF | Hsu et al., 2022[24] | PDB directly (--input-dir) | autoregressive on geometric features (TF32 precision) | --num-samples 1, --seed-per-file --seed-per-sample, --tf32; default sampling temperature | FASTA + summary CSV |
| PiFold | Gao et al., 2023[26] | PDB directory (--pdb_dir) | one-shot parallel decoding | default sampling temperature | FASTA |
| LigandMPNN | Dauparas et al., 2025[27] | PDB sorted by protein-residue length, sharded across 16 parallel workers | autoregressive with explicit ligand/cofactor context | --pack_side_chains 0, --write_pdb 0, --write_packed 0, --split_fasta_subfolders 1; ligand atoms parsed from PDB HETATM | FASTA (one sub-folder per replicate) |
| ABACUS-T | Liu et al., 2024[28] | PDB preprocessed to .npy features by the ABACUS-T preprocessing utility | non-autoregressive discrete diffusion with ESM-2 (650 M) self-conditioning, ligand-aware | --temperature 0.1, --iter_num 30, --PLM_selfcond 1, --selfcondPLM 650M, --augment_eps 0.2, --consider_lig, no MSA prior (--msaprior 0), no fixed positions, chain A | FASTA |
| GraDe-IF (general-protein reference) | Lin et al., 2024[41] | .pt graph (--test_dir) | discrete diffusion (BLOSUM kernel) | --step 100 (DDIM), --diverse, ckpt Sep08_..._63.pt | FASTA |
| EnzymeIF (this work) | this work | same as GraDe-IF | discrete diffusion (BLOSUM kernel) | --step 100 (DDIM), --diverse, ckpt Jul01_..._467.pt | FASTA |
| CatIF (this work) | this work | same as GraDe-IF | discrete diffusion (BLOSUM kernel) | --step 100 (DDIM), --diverse, ckpt Sep24_..._228.pt | FASTA |
| CatIF-RL (this work) | this work | same as GraDe-IF | discrete diffusion (BLOSUM kernel) | --step 100 (DDIM), --diverse, ckpt weight_rl/Jan30_round3/policy_epoch02.pt | FASTA |

#### Common settings (apply to all rows above)

| Item | Value |
|---|---|
| Test backbones | 1,423 enzyme structures (ESMFold-predicted, held-out DLKcat split) |
| Protein chain | A (single chain per backbone) |
| Sequence length cap | ≤ 1,180 AA (matches dataset filter, Table S4) |
| Random seeds | {1111, 2222, 3333, 4444, 5555} |
| Sequences per backbone per method | 5 |

| | |
|---|---|
| Total sequences per method | 1,423 × 5 = 7,115 |

#### Notes

1. **Decoding temperature.** ProteinMPNN, ESM-IF, and PiFold use each method's published default decoding temperature; ABACUS-T uses `temperature 0.1` per its published recommendation for inference. The diffusion-based methods (GraDe-IF / EnzymeIF / CatIF / CatIF-RL) use `--diverse` (categorical sampling at every reverse step) at DDIM `step = 100`. 2. **No MSA / no fixed positions.** Although ABACUS-T supports MSA-prior conditioning and motif-fixing (see manuscript §2.6 for our motif-preserving partial design experiments), the **whole-test-set benchmark in Table 2** uses neither. This isolates each method's structure-only inverse-folding capability. 3. **Ligand context.** LigandMPNN and ABACUS-T explicitly use ligand atoms parsed from PDB HETATM records. The other methods are structure-only and ignore ligands. This is reported faithfully in the "Backbone-conditioning input" column.

**Table S7.** Per-protein metrics on the 1,423-enzyme test set; mean [95 % bootstrap CI] and paired Wilcoxon p (with BH-FDR adj. q) vs CatIF-RL R3 for all four metrics. BH adjustment within each metric (8 baseline comparisons per metric).

| Model | Δlog10 k_cat (mean [95 % CI]) | Δlog10 k_cat p (Wilcoxon) | Δlog10 k_cat q (BH) | Δlog10 k_cat sig | Recovery (mean [95 % CI]) | Recovery p (Wilcoxon) | Recovery q (BH) | Recovery sig | pLDDT (mean [95 % CI]) | pLDDT p (Wilcoxon) | pLDDT q (BH) | pLDDT sig | Backbone RMSD (Å) (mean [95 % CI]) | Backbone RMSD (Å) p (Wilcoxon) | Backbone RMSD (Å) q (BH) | Backbone RMSD (Å) sig |
|---|---|---|---|---|---|---|---|---|---|---|---|---|---|---|---|---|
| **ProteinMPNN** | -0.118 [-0.179, -0.056] | 5.36e-163 | 5.89e-162 | *** | 0.470 [0.467, 0.474] | 3.24e-28 | 5.09e-28 | *** | 86.11 [85.82, 86.39] | 1.56e-137 | 4.30e-137 | *** | 3.860 [3.588, 4.148] | 3.36e-58 | 1.23e-57 | *** |
| **ESM-IF** | 0.075 [0.017, 0.133] | 5.20e-131 | 1.43e-130 | *** | 0.536 [0.530, 0.541] | 0.108 | 0.132 | ns | 82.32 [81.72, 82.90] | 3.11e-67 | 5.70e-67 | *** | 5.199 [4.850, 5.568] | 6.90e-11 | 1.08e-10 | *** |
| **LigandMPNN** | -0.055 [-0.116, 0.008] | 2.57e-145 | 1.41e-144 | *** | 0.514 [0.510, 0.517] | 0.744 | 0.818 | ns | 87.05 [86.73, 87.37] | 1.05e-161 | 1.16e-160 | *** | 3.888 [3.609, 4.183] | 9.48e-63 | 5.21e-62 | *** |
| **PiFold** | -0.086 [-0.148, -0.023] | 2.94e-138 | 1.08e-137 | *** | 0.530 [0.526, 0.534] | 0.011 | 0.015 | * | 85.47 [85.10, 85.84] | 9.91e-138 | 3.63e-137 | *** | 4.228 [3.935, 4.534] | 1.70e-38 | 3.75e-38 | *** |
| **ABACUS-T** | 0.006 [-0.054, 0.0 | 4.22e-120 | 9.28e-120 | *** | 0.604 [0.600, 0.609] | 7.56e-38 | 1.39e-37 | *** | 85.67 [85.38, 85.96] | 1.73e-128 | 3.82e-128 | *** | 4.051 [3.762, 4.356] | 2.97e-55 | 8.16e-55 | *** |

| | | | | | | | | | | | | | | | | |
|---|---|---|---|---|---|---|---|---|---|---|---|---|---|---|---|---|
| | 66] | | | | | | | | | | | | | | | |
| **GraDe-IF** | 0.111 [0.072, 0.153] | 1.24e-99 | 2.27e-99 | *** | 0.462 [0.459, 0.465] | 1.60e-41 | 3.52e-41 | *** | 75.73 [75.18, 76.25] | 8.05e-26 | 1.11e-25 | *** | 5.391 [5.090, 5.702] | 0.011 | 0.013 | * |
| **EnzymeIF** | 0.236 [0.183, 0.289] | 6.19e-87 | 9.73e-87 | *** | 0.709 [0.701, 0.717] | 1.92e-188 | 1.05e-187 | *** | 83.74 [83.28, 84.20] | 3.71e-161 | 2.04e-160 | *** | 3.857 [3.600, 4.119] | 1.01e-68 | 1.11e-67 | *** |
| **CatIF** | 0.365 [0.307, 0.423] | 1.12e-73 | 1.54e-73 | *** | 0.572 [0.562, 0.582] | 2.35e-205 | 2.59e-204 | *** | 79.16 [78.51, 79.81] | 1.13e-45 | 1.78e-45 | *** | 4.885 [4.580, 5.191] | 2.25e-15 | 4.12e-15 | *** |
| **CatIF-RL R1** | 0.406 [0.349, 0.466] | 8.55e-63 | 1.04e-62 | *** | 0.561 [0.551, 0.571] | 9.21e-118 | 3.38e-117 | *** | 78.49 [77.82, 79.16] | 2.59e-11 | 3.17e-11 | *** | 5.092 [4.776, 5.412] | 7.09e-04 | 9.74e-04 | *** |
| **CatIF-RL R2** | 0.519 [0.461, 0.578] | 9.61e-16 | 1.06e-15 | *** | 0.557 [0.548, 0.567] | 9.86e-48 | 2.71e-47 | *** | 77.99 [77.32, 78.67] | 0.808 | 0.888 | ns | 5.177 [4.858, 5.496] | 0.049 | 0.054 | ns |
| **CatIF-RL R3** | 0.605 [0.547, 0.664] | — | — | (reference) | 0.553 [0.544, 0.563] | — | — | (reference) | 77.87 [77.20, 78.56] | — | — | (reference) | 5.322 [4.998, 5.650] | — | — | (reference) |

**Table S8.** CatIF and the three GRPO rounds — per-protein metrics with 95 % CI and joint success rate at multiple δ thresholds.

| Round | Δlog10 k_cat | Recovery | pLDDT | Backbone RMSD (Å) | SR@δ=0.0 | SR@δ=0.5 | SR@δ=1.0 | SR@δ=1.5 |
|---|---|---|---|---|---|---|---|---|
| **CatIF** | 0.365 [0.307, 0.423] | 0.572 [0.562, 0.582] | 79.16 [78.51, 79.81] | 4.885 [4.580, 5.191] | 0.115 | 0.048 | 0.025 | 0.012 |
| **CatIF-RL R1** | 0.406 [0.349, 0.466] | 0.561 [0.551, 0.571] | 78.49 [77.82, 79.16] | 5.092 [4.776, 5.412] | 0.126 | 0.058 | 0.03 | 0.012 |
| **CatIF-RL R2** | 0.519 [0.461, 0.578] | 0.557 [0.548, 0.567] | 77.99 [77.32, 78.67] | 5.177 [4.858, 5.496] | 0.131 | 0.053 | 0.027 | 0.009 |
| **CatIF-RL R3** | 0.605 [0.547, 0.664] | 0.553 [0.544, 0.563] | 77.87 [77.20, 78.56] | 5.322 [4.998, 5.650] | 0.127 | 0.063 | 0.034 | 0.015 |

**Table S9.** Joint success rate sensitivity to the Δlog10 k_cat threshold δ (pLDDT > 90 and backbone RMSD < 4 Å held fixed).

| Model | SR@δ=0.0 | SR@δ=0.25 | SR@δ=0.5 | SR@δ=1.0 | SR@δ=1.5 | SR@δ=2.0 |
|---|---|---|---|---|---|---|
| **ProteinMPNN** | 0.07 | 0.053 | 0.041 | 0.026 | 0.017 | 0.006 |
| **ESM-IF** | 0.094 | 0.077 | 0.056 | 0.027 | 0.015 | 0.007 |
| **LigandMPNN** | 0.11 | 0.088 | 0.069 | 0.034 | 0.02 | 0.009 |
| **PiFold** | 0.089 | 0.068 | 0.048 | 0.031 | 0.016 | 0.007 |
| **ABACUS-T** | 0.065 | 0.048 | 0.035 | 0.02 | 0.012 | 0.006 |
| **GraDe-IF** | 0.014 | 0.007 | 0.004 | 0.001 | 0.001 | 0.001 |
| **EnzymeIF** | 0.125 | 0.087 | 0.06 | 0.027 | 0.013 | 0.007 |
| **CatIF** | 0.115 | 0.077 | 0.048 | 0.025 | 0.012 | 0.003 |
| **CatIF-RL R1** | 0.126 | 0.081 | 0.058 | 0.03 | 0.012 | 0.006 |
| **CatIF-RL R2** | 0.131 | 0.086 | 0.053 | 0.027 | 0.009 | 0.003 |
| **CatIF-RL R3** | 0.127 | 0.085 | 0.063 | 0.034 | 0.015 | 0.007 |

**Table S10.** Pairwise ablation significance — paired Wilcoxon signed-rank tests with Benjamini–Hochberg FDR correction within each metric (6 paired comparisons per metric). Δ(B−A) is the per-protein mean difference (later − earlier stage). Static-pipeline rows test the supervised-stage chain (GraDe-IF → EnzymeIF → CatIF); RL rows test consecutive and overall RL rounds.

| Comparison (A → B) | Δlog10 k_cat Δ(B−A) | Δlog10 k_cat p (Wilcoxon) | Δlog10 k_cat q (BH) | Δlog10 k_cat sig | Recovery Δ(B−A) | Recovery p (Wilcoxon) | Recovery q (BH) | Recovery sig | pLDDT Δ(B−A) | pLDDT p (Wilcoxon) | pLDDT q (BH) | pLDDT sig | Backbone RMSD (Å) Δ(B−A) | Backbone RMSD (Å) p (Wilcoxon) | Backbone RMSD (Å) q (BH) | Backbone RMSD (Å) sig |
|---|---|---|---|---|---|---|---|---|---|---|---|---|---|---|---|---|
| Static: GraDe-IF → EnzymeIF | 0.124 | 4.69e-11 | 5.63e-11 | *** | 0.247 | 5.889999999999999e-234 | 3.5299999999999996e-233 | *** | 8.02 | 4.27e-211 | 2.5600000000000002e-210 | *** | -1.535 | 5.999999999999999e-91 | 3.6e-90 | *** |
| Static: EnzymeIF → CatIF | 0.129 | 1.27e-10 | 1.27e-10 | *** | -0.137 | 1.8100000000000003e-163 | 5.430000000000001e-163 | *** | -4.58 | 6.08e-136 | 1.8299999999999998e-135 | *** | 1.028 | 1.5499999999999998e-40 | 4.6499999999999995e-40 | *** |
| Static: GraDe-IF → CatIF | 0.253 | 6.0700000000000006e-27 | 1.82e-26 | *** | 0.11 | 4.58e-98 | 6.859999999999999e-98 | *** | 3.43 | 5.03e-65 | 1.01e-64 | *** | -0.506 | 4.53e-17 | 9.07e-17 | *** |
| RL: R1 → R2 | 0.112 | 8.959999999999999e-23 | 1.79e-22 | *** | -0.004 | 1.47e-36 | 1.47e-36 | *** | -0.5 | 9.36e-10 | 1.12e-09 | *** | 0.085 | 0.018 | 0.022 | * |
| RL: R2 → R3 | 0.086 | 9.61e-16 | 1.44e-15 | *** | -0.004 | 9.859999999999999e-48 | 1.18e-47 | *** | -0.12 | 0.808 | 0.808 | ns | 0.145 | 0.049 | 0.049 | * |
| RL: R1 → R3 | 0.199 | 8.55e-63 | 5.13e- | *** | -0.008 | 9.21e-118 | 1.84e-117 | *** | -0.62 | 2.59e-11 | 3.89e-11 | *** | 0.23 | 0.000709 | 0.001 | ** |

6
2

**Table S11.** Case-study per-baseline metrics on four representative test enzymes (three global redesigns plus one motif-preserving inpainting). Single-seed showcase values matching manuscript §3.5.

<table>
<tr><th>CASE (EC | PROID)</th><th>LENGTH</th><th>NATIVE KCAT ($S^{-1}$)</th><th>MODEL</th><th>MUTATIONS</th><th>RECOVERY</th><th>ΔLOG10 K_CAT</th><th>BACKBONE RMSD (Å)</th><th>PLDDT</th></tr>
<tr><td>**EC 1.4.1.20 | PROID 570**</td><td>381</td><td>34.0</td><td>ProteinMPNN</td><td>199</td><td>0.478</td><td>-0.26</td><td>3.112</td><td>84.86</td></tr>
<tr><td>**EC 1.4.1.20 | PROID 570**</td><td>381</td><td>34.0</td><td>ESM-IF</td><td>155</td><td>0.593</td><td>-0.201</td><td>4.246</td><td>83.79</td></tr>
<tr><td>**EC 1.4.1.20 | PROID 570**</td><td>381</td><td>34.0</td><td>LigandMPNN</td><td>165</td><td>0.567</td><td>-0.466</td><td>4.069</td><td>86.16</td></tr>
<tr><td>**EC 1.4.1.20 | PROID 570**</td><td>381</td><td>34.0</td><td>PiFold</td><td>170</td><td>0.554</td><td>-0.567</td><td>4.812</td><td>83.92</td></tr>
<tr><td>**EC 1.4.1.20 | PROID 570**</td><td>381</td><td>34.0</td><td>ABACUS-T</td><td>161</td><td>0.577</td><td>-1.067</td><td>3.914</td><td>86.7</td></tr>
<tr><td>**EC 1.4.1.20 | PROID 570**</td><td>381</td><td>34.0</td><td>GraDe-IF</td><td>2</td><td>0.995</td><td>-0.076</td><td>4.216</td><td>84.68</td></tr>
<tr><td>**EC 1.4.1.20 | PROID 570**</td><td>381</td><td>34.0</td><td>EnzymeIF</td><td>139</td><td>0.635</td><td>0.684</td><td>3.394</td><td>85.18</td></tr>
<tr><td>**EC 1.4.1.20 | PROID 570**</td><td>381</td><td>34.0</td><td>CatIF</td><td>144</td><td>0.622</td><td>0.172</td><td>5.546</td><td>85.57</td></tr>
<tr><td>**EC 1.4.1.20 | PROID 570**</td><td>381</td><td>34.0</td><td>CatIF-RL</td><td>146</td><td>0.617</td><td>0.762</td><td>3.578</td><td>90.33</td></tr>
<tr><td>**EC 2.4.2.1 | PROID 2061**</td><td>289</td><td>8.8</td><td>ProteinMPNN</td><td>151</td><td>0.478</td><td>-0.53</td><td>0.696</td><td>86.69</td></tr>
<tr><td>**EC 2.4.2.1 | PROID 2061**</td><td>289</td><td>8.8</td><td>ESM-IF</td><td>105</td><td>0.637</td><td>-0.325</td><td>1.305</td><td>84.73</td></tr>
<tr><td>**EC 2.4.2.1 | PROID 2061**</td><td>289</td><td>8.8</td><td>LigandMPNN</td><td>146</td><td>0.495</td><td>-0.594</td><td>0.946</td><td>85.86</td></tr>
<tr><td>**EC 2.4.2.1 | PROID 2061**</td><td>289</td><td>8.8</td><td>PiFold</td><td>136</td><td>0.529</td><td>-0.059</td><td>0.952</td><td>93.0</td></tr>
<tr><td>**EC 2.4.2.1 | PROID 2061**</td><td>289</td><td>8.8</td><td>ABACUS-T</td><td>104</td><td>0.64</td><td>-0.128</td><td>0.829</td><td>86.6</td></tr>
<tr><td>**EC 2.4.2.1 | PROID 2061**</td><td>289</td><td>8.8</td><td>GraDe-IF</td><td>0</td><td>1.0</td><td>0.0</td><td>2.039</td><td>79.27</td></tr>
<tr><td>**EC 2.4.2.1 | PROID 2061**</td><td>289</td><td>8.8</td><td>EnzymeIF</td><td>0</td><td>1.0</td><td>0.0</td><td>0.002</td><td>92.54</td></tr>
<tr><td>**EC 2.4.2.1 | PROID 2061**</td><td>289</td><td>8.8</td><td>CatIF</td><td>0</td><td>1.0</td><td>0.0</td><td>0.002</td><td>92.54</td></tr>
<tr><td>**EC 2.4.2.1 | PROID 2061**</td><td>289</td><td>8.8</td><td>CatIF-RL</td><td>2</td><td>0.993</td><td>0.227</td><td>0.03</td><td>92.54</td></tr>
<tr><td>**EC 5.3.1.1 | PROID 6002**</td><td>250</td><td>63.0</td><td>ProteinMPNN</td><td>134</td><td>0.464</td><td>-1.428</td><td>0.775</td><td>85.65</td></tr>
<tr><td>**EC 5.3.1.1 | PROID 6002**</td><td>250</td><td>63.0</td><td>ESM-IF</td><td>84</td><td>0.664</td><td>-0.453</td><td>0.545</td><td>89.66</td></tr>
<tr><td>**EC 5.3.1.1 | PROID 6002**</td><td>250</td><td>63.0</td><td>LigandMPNN</td><td>125</td><td>0.5</td><td>-0.972</td><td>0.599</td><td>87.56</td></tr>
<tr><td>**EC 5.3.1.1 | PROID 6002**</td><td>250</td><td>63.0</td><td>PiFold</td><td>123</td><td>0.508</td><td>-0.951</td><td>1.722</td><td>87.18</td></tr>
</table>

| | | | | | | | | |
|---|---|---|---|---|---|---|---|---|
| **EC 5.3.1.1 \| PROID 6002** | 250 | 63.0 | ABACUS-T | 88 | 0.648 | -0.388 | 0.654 | 91.35 |
| **EC 5.3.1.1 \| PROID 6002** | 250 | 63.0 | GraDe-IF | 1 | 0.996 | -0.05 | 1.341 | 80.22 |
| **EC 5.3.1.1 \| PROID 6002** | 250 | 63.0 | EnzymeIF | 6 | 0.976 | -0.214 | 0.053 | 96.19 |
| **EC 5.3.1.1 \| PROID 6002** | 250 | 63.0 | CatIF | 1 | 0.996 | 0.151 | 0.085 | 95.87 |
| **EC 5.3.1.1 \| PROID 6002** | 250 | 63.0 | CatIF-RL | 3 | 0.988 | 0.241 | 0.068 | 96.15 |
| **EC 1.1.1.248 (SALR) — MOTIF INPAINTING \| PROID 4771** | 311 | 2.1 | ProteinMPNN | 151 | 0.514 | 0.153 | 0.772 | 88.54 |
| **EC 1.1.1.248 (SALR) — MOTIF INPAINTING \| PROID 4771** | 311 | 2.1 | ESM-IF | 138 | 0.556 | 0.307 | 6.672 | 89.67 |
| **EC 1.1.1.248 (SALR) — MOTIF INPAINTING \| PROID 4771** | 311 | 2.1 | LigandMPNN | 142 | 0.543 | 0.031 | 1.11 | 89.13 |
| **EC 1.1.1.248 (SALR) — MOTIF INPAINTING \| PROID 4771** | 311 | 2.1 | PiFold | 130 | 0.582 | 0.105 | 4.723 | 93.03 |
| **EC 1.1.1.248 (SALR) — MOTIF INPAINTING \| PROID 4771** | 311 | 2.1 | ABACUS-T | 123 | 0.605 | -0.621 | 1.773 | 85.64 |
| **EC 1.1.1.248 (SALR) — MOTIF INPAINTING \| PROID 4771** | 311 | 2.1 | GraDe-IF | 5 | 0.984 | -0.293 | 3.325 | 86.5 |
| **EC 1.1.1.248 (SALR) — MOTIF INPAINTING \| PROID 4771** | 311 | 2.1 | EnzymeIF | 34 | 0.891 | 0.345 | 2.028 | 92.15 |
| **EC 1.1.1.248 (SALR) — MOTIF INPAINTING \| PROID 4771** | 311 | 2.1 | CatIF | 14 | 0.955 | -0.181 | 0.349 | 94.28 |
| **EC 1.1.1.248 (SALR) — MOTIF INPAINTIN** | 311 | 2.1 | CatIF-RL | 18 | 0.942 | 0.392 | 0.666 | 94.01 |

**Table S12.** Fixed (motif-preserving) residues in the SalR (EC 1.1.1.248) subgraph-inpainting case study, per Geissler et al. (2007).

| Position | Functional role | Reference |
|---|---|---|
| **Asn152** | Catalytic — proton transfer | Geissler et al., 2007 |
| **Ser180** | Catalytic — proton transfer | Geissler et al., 2007 |
| **Tyr236** | Catalytic — proton transfer | Geissler et al., 2007 |
| **Lys240** | Catalytic — proton transfer | Geissler et al., 2007 |

#### S3 Supplementary Figures

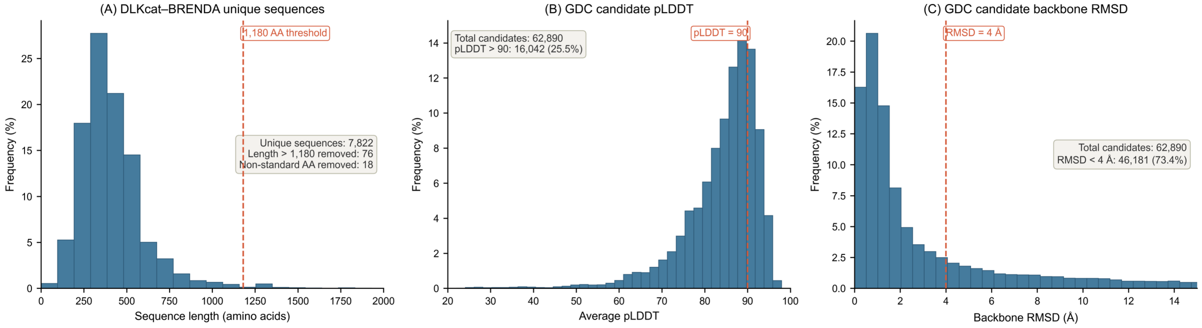

**Figure S1.** Per-stage data filtering distributions across the CatIF-RL data pipeline. (A) DLKcat–BRENDA unique-sequence length distribution with the 1,180 AA cutoff. (B) Average pLDDT distribution of the 62,890 GDC candidate variants with the pLDDT > 90 cutoff. (C) Backbone RMSD distribution with the RMSD < 4 Å cutoff. Pass-count annotations report the number and fraction of candidates satisfying each individual cutoff.

#### (A) EnzymeIF

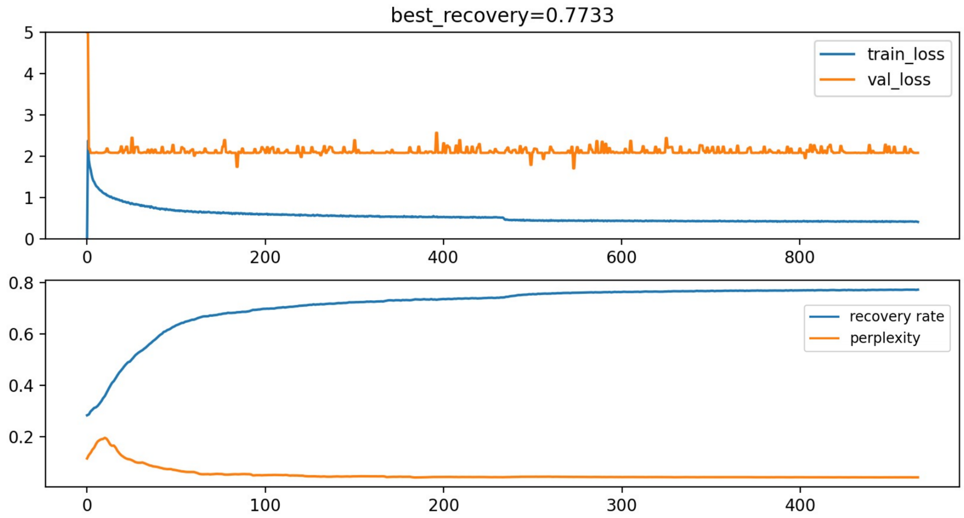

Selected checkpoint: epoch 467 | best validation recovery = 0.7733

#### (A) CatIF

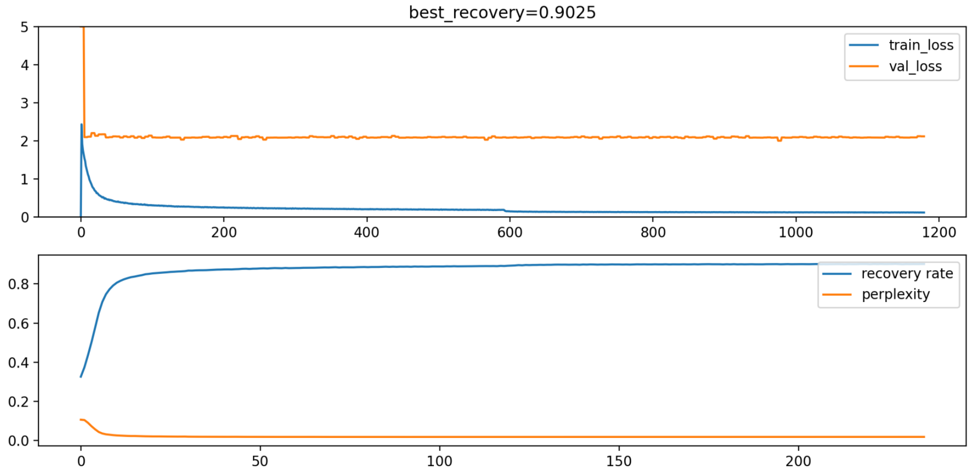

Selected checkpoint: epoch 228 | best validation recovery = 0.9025

**Figure S2.** Training and validation curves for (A) EnzymeIF and (B) CatIF. Top sub-panel of each panel: training loss (blue) and validation loss (orange) versus training step. Bottom sub-panel: validation recovery rate (blue) and scaled perplexity (orange, ×0.01) versus epoch. Selected checkpoints are annotated below each panel.

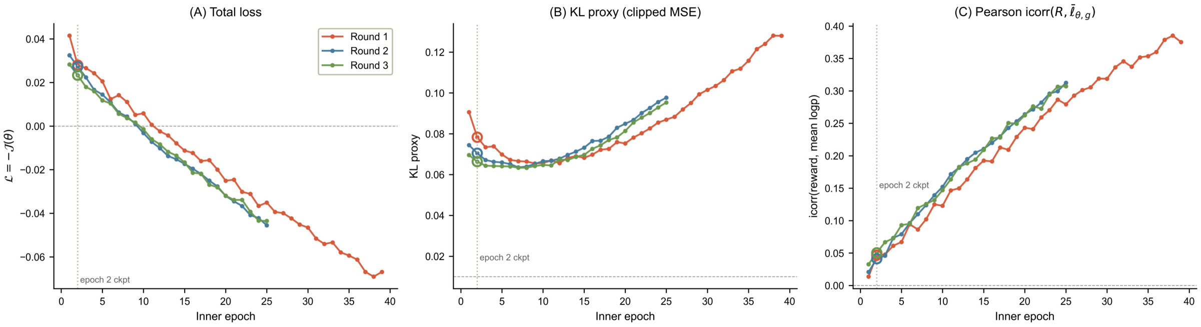

**Figure S3.** CatIF-RL GRPO inner-loop convergence across the three rounds. (A) Total loss = $-\mathcal{J}(\theta)$ per epoch. (B) KL proxy with the KL_target = 0.01 reference dashed line. (C) Pearson correlation between per-sample reward and length-normalised log-probability (icorr(R, ℓ̄_θ,g)). The epoch-2 round-end checkpoint is marked in every panel.

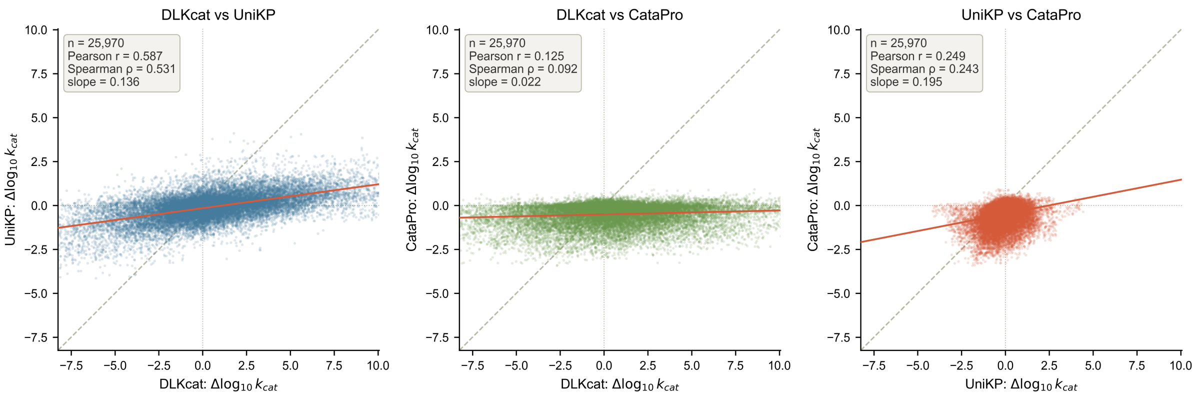

**Figure S4.** Pairwise agreement between the three k_cat predictors (DLKcat, UniKP, CataPro) on 25,970 matched (protein, substrate, mutant) triplets from the EnzymeIF candidate pool. Annotations show Pearson r, Spearman ρ and the linear regression slope. The dashed grey line is y = x; the orange line is the OLS fit.

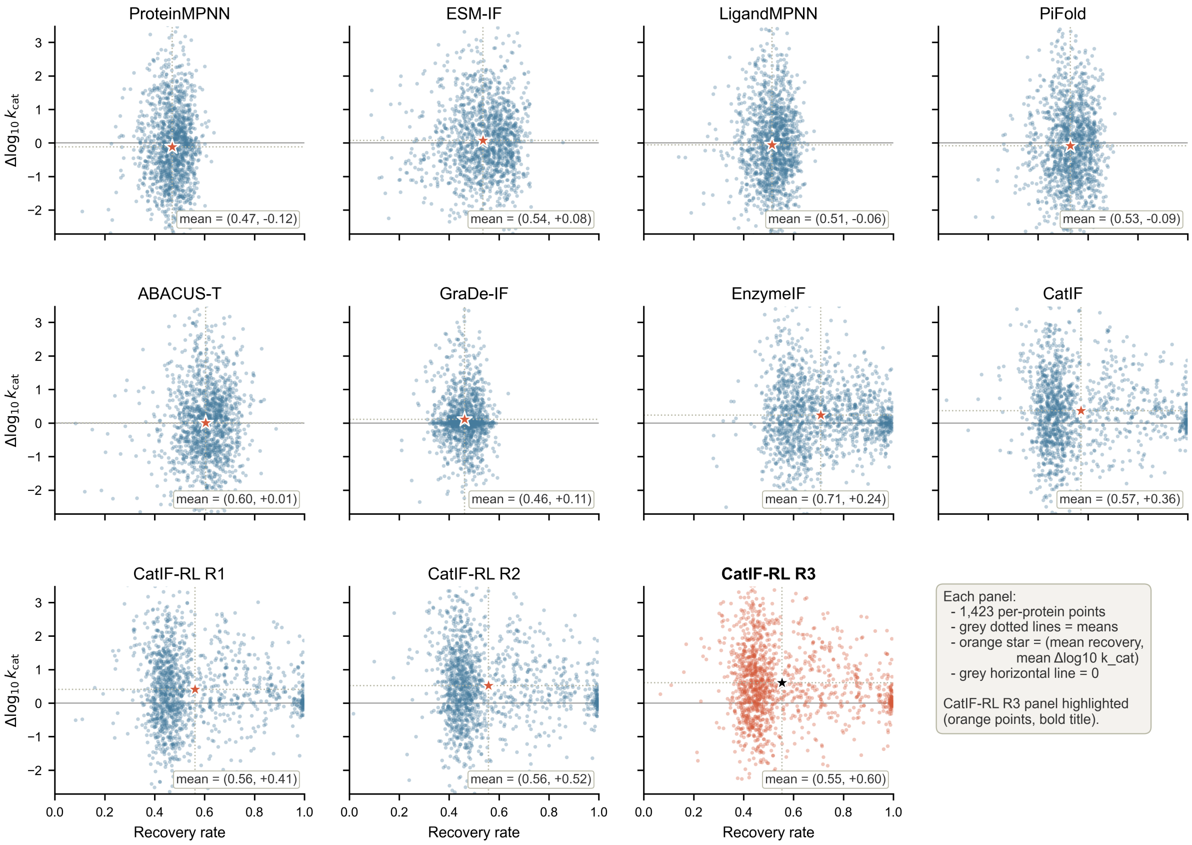

**Figure S5.** Recovery rate versus Δlog10 k_cat per protein, faceted by model. Each panel shows the 1,423 per-protein points for a single model (grey dotted lines mark per-model means, orange star marks the joint mean). CatIF-RL R3 panel highlighted in orange.

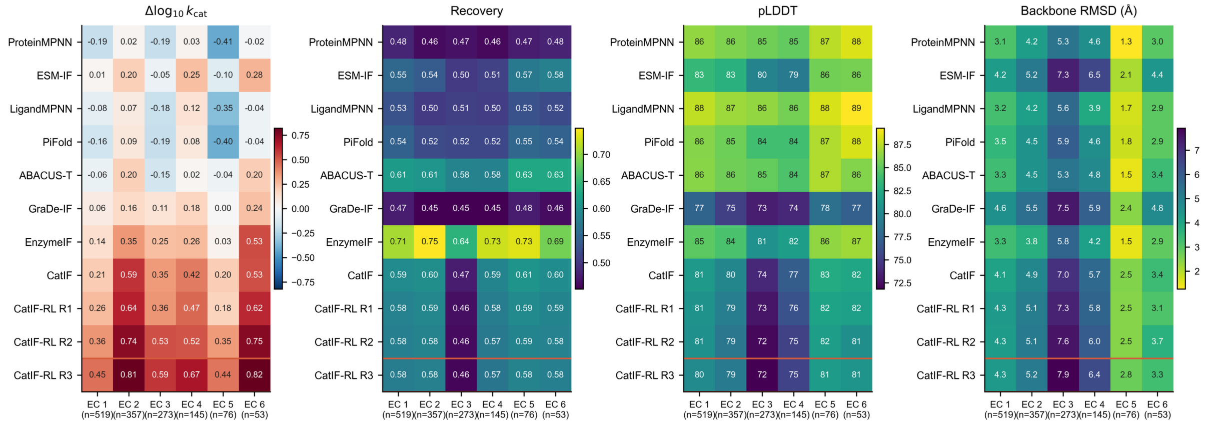

**Figure S6.** EC-class-stratified per-protein performance heatmaps. Rows = models (CatIF-RL R3 highlighted with an orange border), columns = EC main class 1–6 (test-set EC class 7 is empty). Each cell is the per-protein mean of the indicated metric; n per class is shown below the column labels.

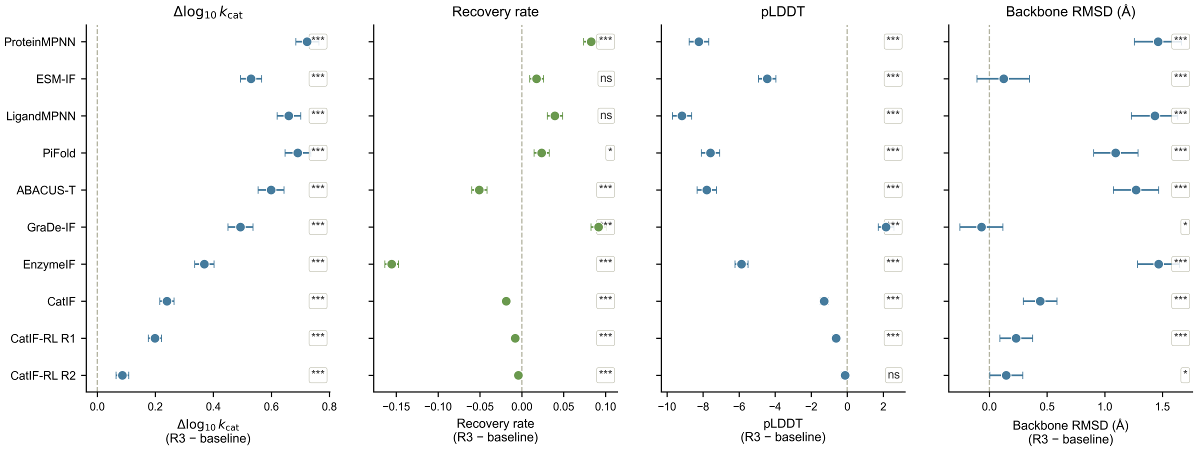

**Figure S7.** Multi-metric significance forest plot. Each panel shows the per-protein mean difference (CatIF-RL R3 − baseline) with 95 % bootstrap confidence interval for every other model on a single metric (Δlog10 k_cat, Recovery, pLDDT, backbone RMSD). Significance stars are BH-FDR-adjusted q-values from paired Wilcoxon signed-rank tests.

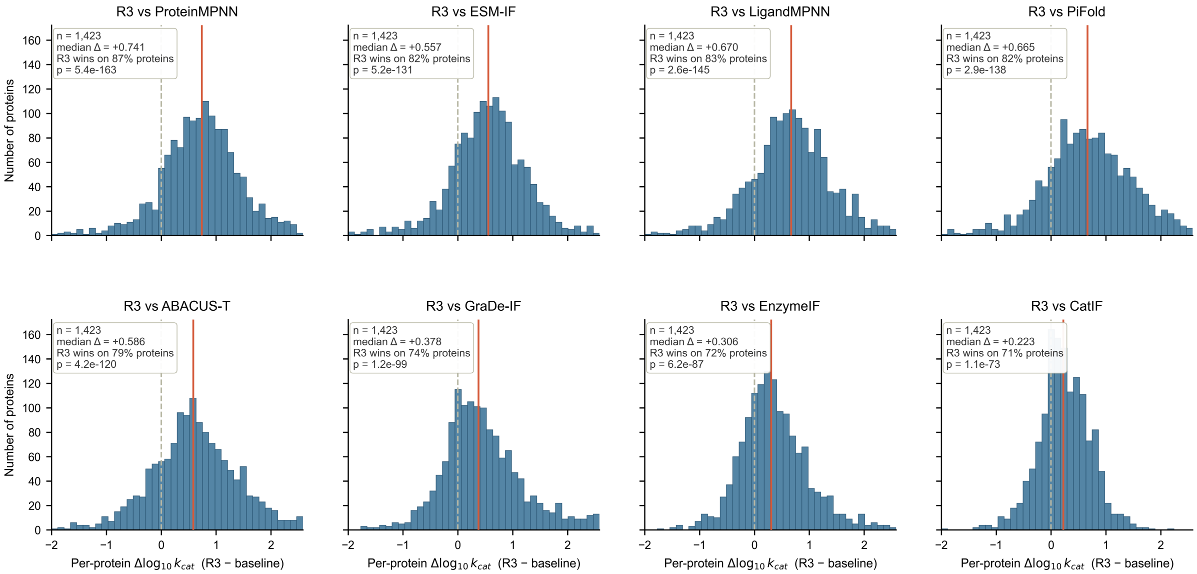

**Figure S8.** Per-protein paired difference histograms (CatIF-RL R3 minus baseline) for Δlog10 k_cat. Each panel shows the distribution of per-protein differences against one baseline. The grey dashed line is zero; the orange line is the median; annotations report n, median, fraction of proteins on which CatIF-RL R3 wins, and the paired Wilcoxon p-value.

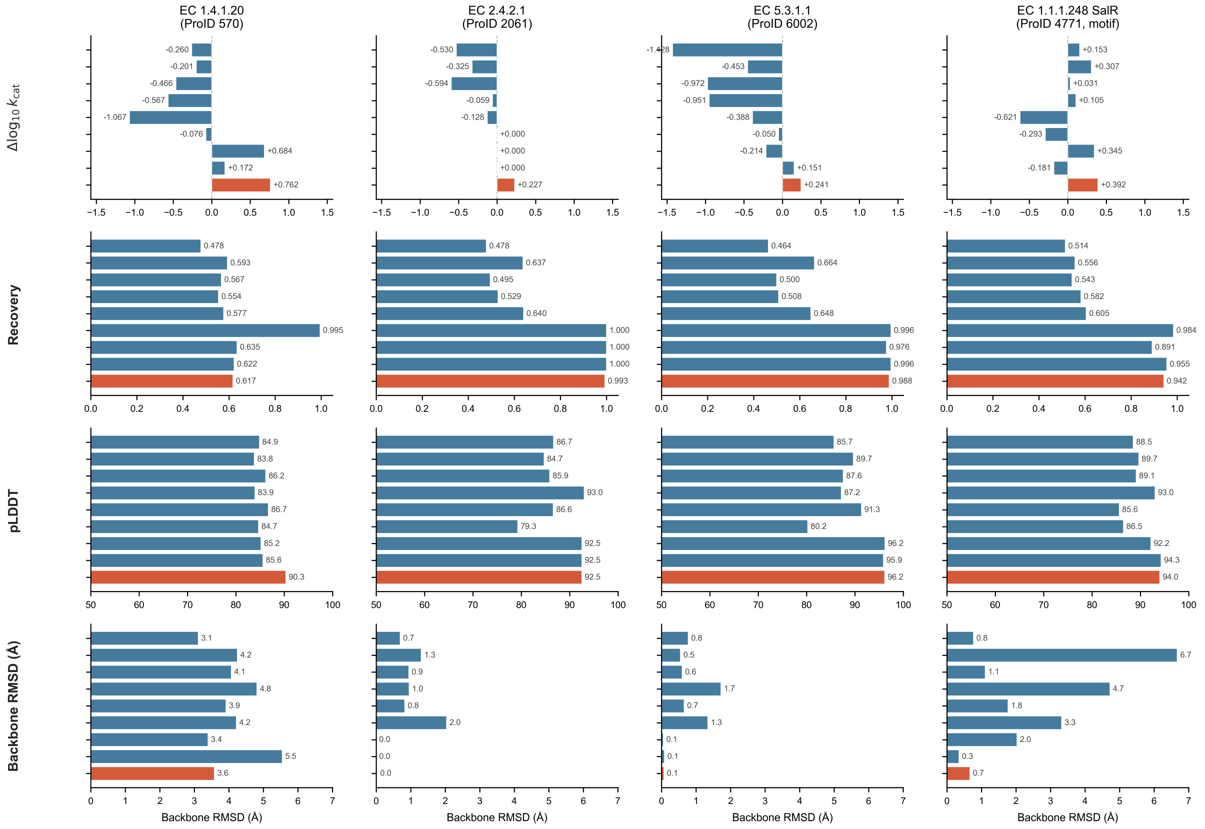

**Figure S9.** Case-study metric grid. Rows = metrics (Δlog10 k_cat, Recovery, pLDDT, backbone RMSD); columns = the four case enzymes. Each cell is a horizontal bar chart of nine baselines for that (case, metric) combination, with CatIF-RL highlighted in orange. Bars are single-seed showcase values matching manuscript §3.5 (seed 1 for CatIF-RL and the seed-numbered baselines; the cumulative-style "12345" file for catif / enzymeif / gradeif). The fourth column (SalR) used motif-preserving inpainting only for CatIF-RL; the other rows in that column were sampled under the same unconstrained protocol as columns 1–3 and are shown for reference only.

### S4 Algorithms

#### Algorithm S1 — Subgraph inpainting on CatIF via RePaint resampling

**Use case.** Given a fixed reference enzyme structure, redesign a user-specified subset of residues (e.g., a binding pocket) while keeping all other residues equal to the wild-type. Used for the four case studies (Section 3.5 of the main text).

**Inputs**

- $\mathcal{G}$ — protein backbone graph (residue nodes + edges)
- `x_native` — ground-truth (native) one-hot residue labels (length N, K = 20)
- $m \in \{0,1\}^N$ — binary mask following manuscript convention: `m_i = 1` denotes a **fixed** position (kept = native), `m_i = 0` denotes a **designable** (redesigned) position
- `p^_θ` — trained discrete diffusion policy (CatIF or CatIF-RL checkpoint)
- `T` — training timesteps (this work: T = 500)
- `Δ` — DDIM stride (this work: Δ = 50)
- `U` — RePaint resampling depth (`jump_n`; this work: **U = 5**)
- `diverse ∈ {True, False}` — stochastic categorical sampling vs. argmax decoding

**Output**

- `x^` — designed sequence (one-hot, length N) consistent with `m`

```
x_T  ← Uniform(K = 20)                                      // initialise noise on all N nodes
for  t = T, T−Δ, T−2Δ, …, Δ  do
    s   ← max(t − Δ, 0)                                     // target step of this DDIM jump
    U_t ← 1  if  s = 0  else  U                             // no jump-back at the final step
    for  u = 1, …, U_t  do
        x_s^var ← Sample( p^_θ(x_s | x_t, 𝒢) )              // marginalised reverse posterior, manuscript §2.6
        x_s^fix ← Categorical( x_native · Q¯_s )            // forward-corrupt native to step s
        x_s     ← m ⊙ x_s^fix + (1 − m) ⊙ x_s^var           // fused state, manuscript §2.6
        if  u < U_t  and  s > 0  then                       // jump back  s → t (re-corrupt)
            Q_{t|s} ← TransitionMatrix( ᾱ_t / ᾱ_s )         // uniform-noise variant only
            x_t ← Categorical( Q_{t|s} · x_s )
        end if
    end for
    x_t ← x_s                                               // advance outer loop
end for
x^ ← argmax_K p^_θ(x_0 | x_Δ, 𝒢)   if  diverse = False
    else  Categorical( p^_θ(x_0 | x_Δ, 𝒢) )
```

```
18: return x^
```

**Notes.** 1. The mask `m` is built from a user-supplied 0-based residue index list (e.g., `"5,10-20,35"`) that selects **designable** positions (`m_i = 0`); all other positions are **fixed** to native (`m_i = 1`). 2. Line 6: `p^_θ(x_s | x_t, 𝒢)` is computed by marginalising over the predicted clean-sequence distribution: `p^_θ(x_s | x_t, 𝒢) = Σ_{k=1}^{K} q(x_s | x_t, x_0=k) · p^_θ(x_0=k | x_t, 𝒢)`, exactly as in manuscript §2.6. 3. Line 7: `Q¯_s = ∏_{i=1}^{s} Q_i` is the cumulative discrete transition matrix (BLOSUM kernel for CatIF). The Categorical samples a noisy version of the native sequence on the correct forward marginal at step `s`. 4. Lines 9–11 (jump-back) admit a closed-form transition `Q_{t|s}` only for the uniform-noise schedule. The case-study runs use **U = 5** with the BLOSUM-noise CatIF policy, where the closed-form jump-back is unavailable; the inner loop therefore reduces to **iterative fusion** — at each outer (t → s) step, the reverse-step → GT-injection operation is repeated U times against the unchanged mask, progressively refining the redesigned region's compatibility with the surrounding native context. 5. The GraDe-IF backbone, `Q¯_t`, the BLOSUM transition kernel, and the marginalised reverse step are reused unchanged from Yi et al. [41]; the iterative-fusion mechanism extends RePaint (Lugmayr et al. [43]) to the BLOSUM-noise discrete-diffusion setting.

#### Algorithm S2 — Iterated KL-regularized offline GRPO refinement of CatIF

**Use case.** Refine a pre-trained CatIF policy $\pi_0$ toward higher predicted Δlogkcat using offline group-relative policy optimization with three sample → score → train rounds.

**Top-level (outer round loop, K = 3):**

```
Input:
  π_θ^(0)     ← pre-trained CatIF (fixed BLOSUM diffusion policy)
  D_pairs     ← (enzyme, substrate) pairs from BRENDA train+valid split
  G           ← group size for sampling per pair (this work: G = 5)
  K           ← number of outer rounds (this work: K = 3)
  Φ           ← predictor ensemble {DLKcat, UniKP, CataPro} → S_ensemble

Output:
  π_θ^(K)     ← final CatIF-RL policy

π_ref ← π_θ^(0)                                          // frozen reference for KL anchor (manuscript §2.5)
for k = 1, …, K do
    Σ_k ← { sample G candidates per (enzyme, substrate) using π_θ^(k−1); DDIM step = 100 }
    For each candidate, compute S_ensemble via Φ              // manuscript §2.3, §2.5
    D_k ← {(enzyme, substrate, σ, S_ensemble)}                // offline-scored CSV
    π_θ^(k) ← GRPO_train(π_θ^(k−1), π_ref, D_k; E = 2)        // inner loop, see below; epoch-02 ckpt selected
end for
return π_θ^(K)
```

**Inner loop (GRPO_train) — called at each round k**

**Hyperparameters (see Table S3 for values).** epochs `E`, learning rate η, weight decay `wd`, KL target `KL_target`, KL clip `c`, max `β_max`, min `β_min = 5e-4`, gradient clip `clip_g`, gradient accumulation `accum`, sampling step (used for log-prob evaluation), reward shaping params (`τ`, `λ_mut`, $\mu_0$), group filter threshold `R_min = 3`, group size `G` (inherited from outer loop).

```
Input:
  π_θ       ← initial policy this round (= π_θ^(k-1))
  π_ref     ← frozen reference (= π_θ^(0), original CatIF)
  D_k       ← offline-scored dataset for round k

Group D_k by (ProID, SMILES); within each group dedup by sequence.
Drop groups with fewer than R_min distinct rewards.            // ensures learnable signal
β ← β_0
for epoch = 1, …, E  do
    Shuffle the surviving groups.
    for each group g  do
        ## Build inputs
        batch ← G clones of the conditioning graph 𝒢_g
        x_mut ← inject(σ_g,i for i = 1..G)                        // one-hot mutant residues
        μ_g    ← mutation_fraction(x_native_g, x_mut)             // per-sample vector ∈ [0,1]^G

        ## Reward shaping (manuscript §2.5)
        r_act ← clip( S_ensemble_g / τ, −1, 1 )                   // this work: lin_sym
        r_mut ← − λ_mut · max( 0, μ_g − μ_0 )                     // mutation-budget penalty
        R     ← r_act + r_mut                                     // total reward

        ## Group-relative advantage
        A     ← (R − R̄_g) / (σ_g + ε)

        ## Length-normalized policy log-prob (manuscript §2.5)
        ℓ̄_θ,g    ← (1 / L_g) · Σ_t  log π_θ( x_{g,t} | 𝒢_g )      // requires_grad = True
        ℓ̄_ref,g  ← (1 / L_g) · Σ_t  log π_ref( x_{g,t} | 𝒢_g )    // no_grad

        ## KL proxy (manuscript Eq. for 𝒥(θ))
        κ        ← clip( ℓ̄_θ,g − ℓ̄_ref,g, −c, c )
        KL_proxy ← (1 / G)  · Σ_g  κ²

        ## Loss  L = −𝒥(θ)
        L_pg     ← −(1 / G)  · Σ_g  stop_grad(A)  · ℓ̄_θ,g
        L        ← (L_pg + β  · KL_proxy) / accum
        backward(L)
        if accumulated_steps % accum = 0:
            clip_grad_norm_(π_θ, clip_g)
            optimizer.step();  optimizer.zero_grad()

```

```
        ## Adaptive β
        if   KL_proxy > 1.5 · KL_target :  β ← min( β · 1.5, β_max )
        elif KL_proxy < 0.5 · KL_target :  β ← max( β · 0.7, β_min )
    end for
    save_checkpoint(π_θ, epoch)
end for
return π_θ                                              // epoch-02 checkpoint used downstream
```

**Notes.** 1. **Offline GRPO.** Rewards (S_ensemble) are pre-computed by the predictor ensemble and stored in CSV; the trainer never queries the predictors during gradient updates. This avoids the cost and stochasticity of online predictor calls. 2. **Length-normalised log-prob `ℓ_θ,g`.** Following manuscript §2.5, log-probabilities are token-averaged over the sequence length L_g. This prevents long sequences from dominating the gradient — a length-normalised analogue of GRPO's per-token credit assignment. 3. **Filter `R_min ≥ 3`.** Groups whose rewards collapse to one or two distinct values produce trivial advantages, so they are excluded from the gradient pool. 4. **KL proxy.** The exact KL between two multinomial diffusion policies is intractable; we substitute a clipped MSE on the token-mean log-prob gap (manuscript Eq. for $\mathcal{J}(\theta)$), which is differentiable and well-behaved under AMP/fp16. 5. **Adaptive β.** β is updated multiplicatively after every group, bounded in [β_min, β_max]. This keeps the policy from drifting too far while still permitting useful gradient flow when KL is well below target. 6. **Reward shaping.** This work uses lin_sym: r_act = clip(S_ensemble / τ, ±1), with τ = 0.40. The mutation-budget penalty λ_mut · max(0, μ − $\mu_0$) activates only when the per-sample mutation fraction exceeds $\mu_0$ = 0.30, encouraging conservative edits. 7. **Reference policy.** Across all K = 3 rounds, π_ref is held fixed at the original CatIF checkpoint π_θ^(0) (not the previous round's policy). This anchors the KL term to a stable baseline and prevents reward hacking. 8. **Operational E = 2.** Each round's training is capped at the epoch-02 checkpoint to mitigate over-fitting on the fixed offline batch (see Table S3 note 1).

(46)Guo, D.; Yang, D.; Zhang, H.; Song, J.; Wang, P.; Zhu, Q.; Xu, R.; Zhang, R.; Ma, S.; Bi, X.; Zhang, X.; Yu, X.; Wu, Y.; Wu, Z. F.; Gou, Z.; Shao, Z.; Li, Z.; Gao, Z.; Liu, A.; Xue, B.; Wang, B.; Wu, B.; Feng, B.; Lu, C.; Zhao, C.; Deng, C.; Ruan, C.; Dai, D.; Chen, D.; Ji, D.; Li, E.; Lin, F.; Dai, F.; Luo, F.; Hao, G.; Chen, G.; Li, G.; Zhang, H.; Xu, H.; Ding, H.; Gao, H.; Qu, H.; Li, H.; Guo, J.; Li, J.; Chen, J.; Yuan, J.; Tu, J.; Qiu, J.; Li, J.; Cai, J. L.; Ni, J.; Liang, J.; Chen, J.; Dong, K.; Hu, K.; You, K.; Gao, K.; Guan, K.; Huang, K.; Yu, K.; Wang, L.; Zhang, L.; Zhao, L.; Wang, L.; Zhang, L.; Xu, L.; Xia, L.; Zhang, M.; Zhang, M.; Tang, M.; Zhou, M.; Li, M.; Wang, M.; Li, M.; Tian, N.; Huang, P.; Zhang, P.; Wang, Q.; Chen, Q.; Du, Q.; Ge, R.; Zhang, R.; Pan, R.; Wang, R.; Chen, R. J.; Jin, R. L.; Chen, R.; Lu, S.; Zhou, S.; Chen, S.; Ye, S.; Wang, S.; Yu, S.; Zhou, S.; Pan, S.; Li, S. S.; Zhou, S.; Wu, S.; Yun, T.; Pei, T.; Sun, T.; Wang, T.; Zeng, W.; Liu, W.; Liang, W.; Gao, W.; Yu, W.; Zhang, W.; Xiao, W. L.; An, W.; Liu, X.; Wang, X.; Chen, X.; Nie, X.; Cheng, X.; Liu, X.; Xie, X.; Liu, X.; Yang, X.; Li, X.; Su, X.; Lin, X.; Li, X. Q.; Jin, X.; Shen, X.; Chen, X.; Sun, X.; Wang, X.; Song, X.; Zhou, X.; Wang, X.; Shan, X.; Li, Y. K.; Wang, Y. Q.; Wei, Y. X.; Zhang, Y.; Xu, Y.; Li, Y.; Zhao, Y.; Sun, Y.; Wang, Y.; Yu, Y.; Zhang, Y.; Shi, Y.; Xiong, Y.; He, Y.; Piao, Y.; Wang, Y.; Tan, Y.; Ma, Y.; Liu, Y.; Guo, Y.; Ou, Y.; Wang, Y.; Gong, Y.; Zou, Y.; He, Y.; Xiong, Y.; Luo, Y.; You, Y.; Liu, Y.; Zhou, Y.; Zhu, Y. X.; Huang, Y.; Li, Y.; Zheng, Y.; Zhu, Y.; Ma, Y.; Tang, Y.; Zha, Y.; Yan, Y.; Ren, Z. Z.; Ren, Z.; Sha, Z.; Fu, Z.; Xu, Z.; Xie, Z.; Zhang, Z.; Hao, Z.; Ma, Z.; Yan, Z.; Wu, Z.; Gu, Z.; Zhu, Z.; Liu, Z.; Li, Z.; Xie, Z.; Song, Z.; Pan, Z.; Huang, Z.; Xu, Z.; Zhang, Z.; Zhang, Z. DeepSeek-R1 Incentivizes Reasoning in LLMs through Reinforcement Learning. *Nature* **2025**, *645* (8081), 633–638. https://doi.org/10.1038/s41586-025-09422-z.

(47)Lugmayr, A.; Danelljan, M.; Romero, A.; Yu, F.; Timofte, R.; Gool, L. V. RePaint: Inpainting Using Denoising Diffusion Probabilistic Models; IEEE Computer Society, 2022; pp 11451–11461. https://doi.org/10.1109/CVPR52688.2022.01117.

(48)Lin, Z.; Akin, H.; Rao, R.; Hie, B.; Zhu, Z.; Lu, W.; Smetanin, N.; Verkuil, R.; Kabeli, O.; Shmueli, Y.; dos Santos Costa, A.; Fazel-Zarandi, M.; Sercu, T.; Candido, S.; Rives, A. Evolutionary-Scale Prediction of Atomic-Level Protein Structure with a Language Model. *Science* **2023**, *379* (6637), 1123–1130. https://doi.org/10.1126/science.ade2574.

(49)Geissler, R.; Brandt, W.; Ziegler, J. Molecular Modeling and Site-Directed Mutagenesis Reveal the Benzylisoquinoline Binding Site of the Short-Chain Dehydrogenase/Reductase Salutaridine Reductase. *Plant Physiol* **2007**, *143* (4), 1493–1503. https://doi.org/10.1104/pp.106.095166.

(50)Luo, S.; Su, Y.; Peng, X.; Wang, S.; Peng, J.; Ma, J. Antigen-Specific Antibody Design and Optimization with Diffusion-Based Generative Models for Protein Structures. In *Advances in Neural Information Processing Systems*; 2022; Vol. 35, pp 9754–9767.

(51)Wang, J.; Lisanza, S.; Juergens, D.; Tischer, D.; Watson, J. L.; Castro, K. M.; Ragotte, R.; Saragovi, A.; Milles, L. F.; Baek, M.; Anishchenko, I.; Yang, W.; Hicks, D. R.; Expòsit, M.; Schlichthaerle, T.; Chun, J.-H.; Dauparas, J.; Bennett, N.; Wicky, B. I. M.; Muenks, A.; DiMaio, F.; Correia, B.; Ovchinnikov, S.; Baker, D. Scaffolding Protein Functional Sites Using Deep Learning. *Science* **2022**, *377* (6604), 387–394. https://doi.org/10.1126/science.abn2100.

(52)Sillitoe, I.; Bordin, N.; Dawson, N.; Waman, V. P.; Ashford, P.; Scholes, H. M.; Pang, C. S. M.; Woodridge, L.; Rauer, C.; Sen, N.; Abbasian, M.; Le Cornu, S.; Lam, S. D.; Berka, K.; Varekova, I. H.; Svobodova, R.; Lees, J.; Orengo, C. A. CATH: Increased Structural Coverage of Functional Space. *Nucleic Acids Res* **2021**, *49* (D1), D266–D273. https://doi.org/10.1093/nar/gkaa1079.

(53)Gasser, H.-C.; Oyarzún, D. A.; Alfaro, J. A.; Rajan, A. Tuning ProteinMPNN to Reduce Protein Visibility via MHC Class I through Direct Preference Optimization.
